# Chromosome contacts in activated T cells identify autoimmune disease candidate genes

**DOI:** 10.1101/100958

**Authors:** Oliver S Burren, Arcadio Rubio García, Biola-Maria Javierre, Daniel B Rainbow, Jonathan Cairns, Nicholas J Cooper, John J Lambourne, Ellen Schofield, Xaquin Castro Dopico, Ricardo C Ferreira, Richard Coulson, Frances Burden, Sophia P Rowlston, Kate Downes, Steven W Wingett, Mattia Frontini, Willem H Ouwehand, Peter Fraser, Mikhail Spivakov, John A Todd, Linda S Wicker, Antony J Cutler, Chris Wallace

**Affiliations:** Department of Medicine, University of Cambridge, Addenbrooke’s Hospital, Cambridge, CB2 0SP, UK; JDRF/Wellcome Trust Diabetes and Inflammation Laboratory, Department of Medical Genetics, NIHR Cambridge Biomedical Research Centre, Cambridge Institute for Medical Research, University of Cambridge, Cambridge, CB2 0XY, UK; Current address: JDRF/Wellcome Trust Diabetes and Inflammation Laboratory, Wellcome Trust Centre for Human Genetics, Nuffield Department of Medicine, NIHR Oxford Biomedical Research Centre, University of Oxford, Roosevelt Drive, Oxford, OX3 7BN, UK; Nuclear Dynamics Programme, The Babraham Institute, Babraham Research Campus, Cambridge, CB22 3AT, UK; Department of Haematology, University of Cambridge, Cambridge Biomedical Campus, Long Road, Cambridge, CB2 0PT, UK; National Health Service Blood and Transplant, Cambridge Biomedical Campus, Long Road, Cambridge, CB2 0PT, UK; British Heart Foundation Centre of Excellence, Division of Cardiovascular Medicine, Addenbrooke’s Hospital, Hills Road, Cambridge, CB2 0QQ, UK; Department of Human Genetics, Wellcome Trust Sanger Institute, Wellcome Trust Genome Campus, Hinxton, Cambridge, CB10 1HH, UK; MRC Biostatistics Unit, University of Cambridge, Cambridge Institute of Public Health, Cambridge Biomedical Campus, Cambridge, CB2 0SR, UK

**Keywords:** genetics, genomics, chromatin conformation, CD4^+^ T cells, CD4^+^ T cell activation, autoimmune disease, genome-wide association studies

## Abstract

**Background:** Autoimmune disease-associated variants are preferentially found in regulatory regions in immune cells, particularly CD4^+^ T cells. Linking such regulatory regions to gene promoters in disease-relevant cell contexts facilitates identification of candidate disease genes.

**Results:** Within four hours, activation of CD4^+^ T cells invokes changes in histone modifications and enhancer RNA transcription that correspond to altered expression of the interacting genes identified by promoter capture Hi-C. By integrating promoter capture Hi-C data with genetic associations for five autoimmune diseases we prioritised 245 candidate genes with a median distance from peak signal to prioritised gene of 153 kb. Just under half (108/245) prioritised genes related to activation-sensitive interactions. This included *IL2RA*, where allele-specific expression analyses were consistent with its interaction-mediated regulation, illustrating the utility of the approach.

**Conclusions:** Our systematic experimental framework offers an alternative approach to candidate causal gene identification for variants with cell state-specific functional effects, with achievable sample sizes.

## Background

Genome-wide association studies (GWAS) in the last decade have associated 324 distinct genomic regions to at least one and often several autoimmune diseases (http://www.immunobase.org). The majority of associated variants lie outside genes[1] and presumably tag regulatory variants acting on nearby or more distant genes[2, 3]. Progress from GWAS discovery to biological interpretation has been hampered by lack of systematic methods to define the gene(s) regulated by a given variant. The use of Hi-C[4] and capture Hi-C to link GWAS identified variants to their target genes for breast cancer[5] and autoimmune diseases[6] using cell lines, has highlighted the potential for mapping long range interactions in advancing our understanding of disease association. The observed cell specificity of these interactions indicates a need to study primary disease-relevant human cells, and investigate the extent to which cell state may affect inference.

Integration of GWAS signals with cell-specific chromatin marks has highlighted the role of regulatory variation in immune cells[7], and in particular CD4^+^ T cells, in autoimmune diseases[8]. Concordantly, differences in DNA methylation of immune-related genes have been observed in CD4^+^ T cells from autoimmune disease patients compared to healthy controls [9, 10]. CD4^+^ T cells are at the centre of the adaptive immune system and exquisite control of activation is required to guide CD4^+^ T cell fate through selection, expansion and differentiation into one of a number of specialised subsets. Additionally, the prominence of variants in physical proximity to genes associated with T cell regulation in autoimmune disease GWAS and the association of human leukocyte antigen haplotypes have suggested that control of T cell activation is a key etiological pathway in development of many autoimmune diseases[11].

We explored the effect of activation on CD4^+^ T cell gene expression, chromatin states and chromosome conformation. Promoter capture Hi-C (PCHi-C) was used to map promoter interacting regions (PIRs), and to relate activation-induced changes in gene expression to changes in chromosome conformation and transcription of PCHi-C linked enhancer RNAs (eRNAs). We also fine mapped the most probable causal variants for five autoimmune diseases, autoimmune thyroid disease (ATD), coeliac disease (CEL), rheumatoid arthritis (RA), systemic lupus erythematosus (SLE) and type 1 diabetes (T1D). We integrated these sources of information to derive a systematic prioritisation of candidate autoimmune disease genes.

## Results

### A time-course expression profile of early CD4^+^ T cell activation

We profiled gene expression in CD4^+^ T cells from 20 healthy individuals across a 21 hour activation time-course, and identified eight distinct gene modules by clustering these profiles (Fig. 1, Additional File 1: Table S1). This experimental approach focused on much earlier events than previous large time-course studies (eg 6 hours - 8 days[12]) and highlights the earliest changes that are either not seen or are returning towards baseline by 6 hours (Additional File 2: Fig. S1). Gene set enrichment analysis using MSigDB Hallmark gene sets[13] demonstrated that these modules captured temporally distinct aspects of CD4^+^ T cell activation. For example, negative regulators of TGF-beta signalling were rapidly upregulated, and returned to baseline by 4 hours. Interferon responses, inflammatory responses and IL-2 and STAT5 signalling pathways showed a more sustained upregulation out beyond 6 hours, while fatty acid metabolism was initially downregulated in favour of oxidative phosphorylation.

**Fig. 1.**
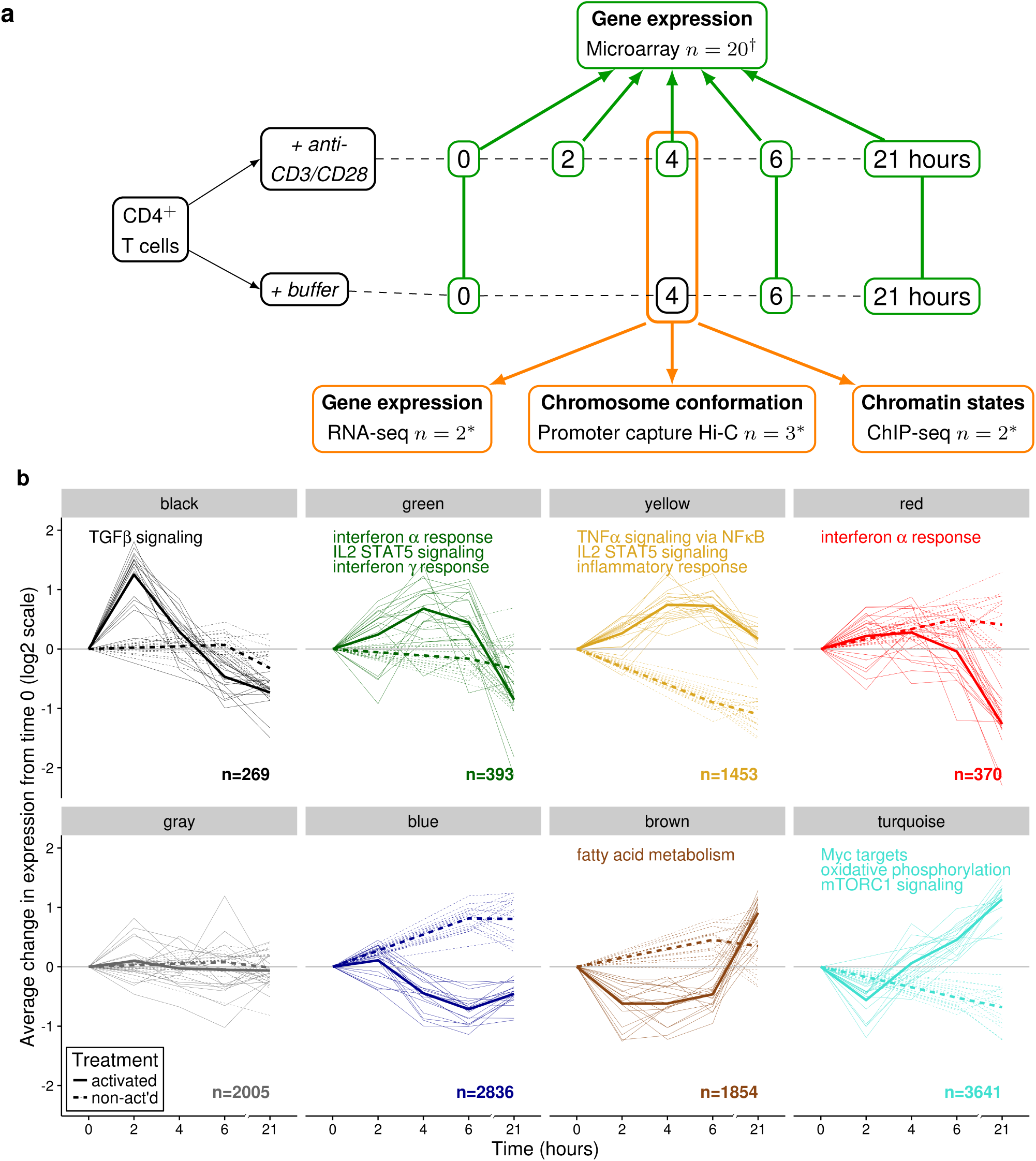
**a** Summary of genomic profiling of CD4^+^ T cells during activation with anti-CD3/CD28 beads. We examined gene expression using microarray in activated and non-activated CD4^+^ T cells across 21 hours, and assayed cells in more detail at the four hour time point using ChIP-seq, RNA-seq and PCHi-C. n gives the number of individuals† or pools* assayed. **b** Eight modules of co-regulated genes were identified, and eigengenes are plotted for each individual (solid lines=activated, dashed lines=non-activated), with heavy lines showing the average eigengene across individuals. We characterized these modules by gene set enrichment analysis within the MSigDB HALLMARK gene sets, and where significant gene sets were found, up to three are shown per module. n is the number of genes in each module.

### PCHi-C captures dynamic enhancer-promoter interactions

We examined activated and non-activated CD4^+^ T cells purified from healthy individuals in more detail at the four-hour time point, at which the average fold change of genes related to cytokine signalling and inflammatory response was maximal, using total RNA sequencing, histone mark chromatin immunoprecipitation sequencing (ChIP-seq) and PCHi-C. Of 8,856 genes identified as expressed (see Methods) in either condition (non-activated or activated), 25% were up- or down-regulated (1,235 and 952 genes respectively, FDR<0.01, Additional File 3: Table S2). We used PCHi-C to characterise promoter interactions in activated and non-activated CD4^+^ T cells. Our capture design baited 22,076 *Hind*III fragments (median length 4 kb) which contained the promoters of 29,131 annotated genes, 18,202 of which are protein coding (Additional File 4: Table S3). We detected 283,657 unique PCHi-C interactions with the CHiCAGO pipeline[14], with 55% found in both activation states, and 22% and 23% found only in non-activated and only in activated CD4^+^ T cells, respectively (Additional File 5: Table S4). 11,817 baited promoter fragments were involved in at least one interaction, with a median distance between interacting fragments of 324 kb. Each interacting promoter fragment connected to a median of eight promoter interacting regions (PIRs), while each interacting PIR was connected to a median of one promoter fragment (Additional File 2: Fig. S2).

We compared our interaction calls to an earlier ChIA-PET dataset from non-activated CD4^+^ T cells[15] and found we replicated over 50% of the longer range interactions (100 kb or greater), with replication rates over ten-fold greater for interactions found in non-activated CD4^+^ T cells compared to interactions found only in erythroblasts or megakaryocytes (Additional File 2: Fig. S3). We also compared histone modification profiles in interacting fragments in CD4^+^ T cells to interacting fragments found in erythroblasts or megakaryocytes. Both promoter fragments and, to a lesser extent, PIRs were enriched for histone modifications associated with transcriptionally active genes and regulatory elements (H3K27ac, H3K4me1, H3K4me3; Additional File 2: Fig. S4), and changes in H3K27ac modifications at both promoter fragments and PIRs correlated with changes in gene expression upon activation. PIRs, but not promoter fragments, showed significant overlap with regions previously annotated as enhancers[16].

We found that absolute levels of gene expression correlated with the number of PIRs (Additional File 2: Fig. S5a, rho=0.81), consistent with recent observations[15]. We defined a subset of PCHi-C interactions that were specifically gained or lost upon activation (2,334 and 1,866 respectively, FDR<0.01) and found that the direction of change (gain or loss) at these differential interactions agreed with the direction of differential expression (up- or down-regulated) at the module level (**Fig. 2**). We further found that dynamic changes in gene expression upon activation correlated with changing numbers of PIRs. Notably, the pattern was asymmetrical, with a gained interaction associating with approximately two fold the change associated with a lost interaction (**Fig. 3a**). Given the >6 hour median half life of mRNAs expressed in T cells[17] (Additional File 2: Fig. S5b), it is possible that the relatively smaller changes associated with lost interactions are due to the persistence of downregulated transcripts at the early stages of T cell activation.

**Fig. 2.**
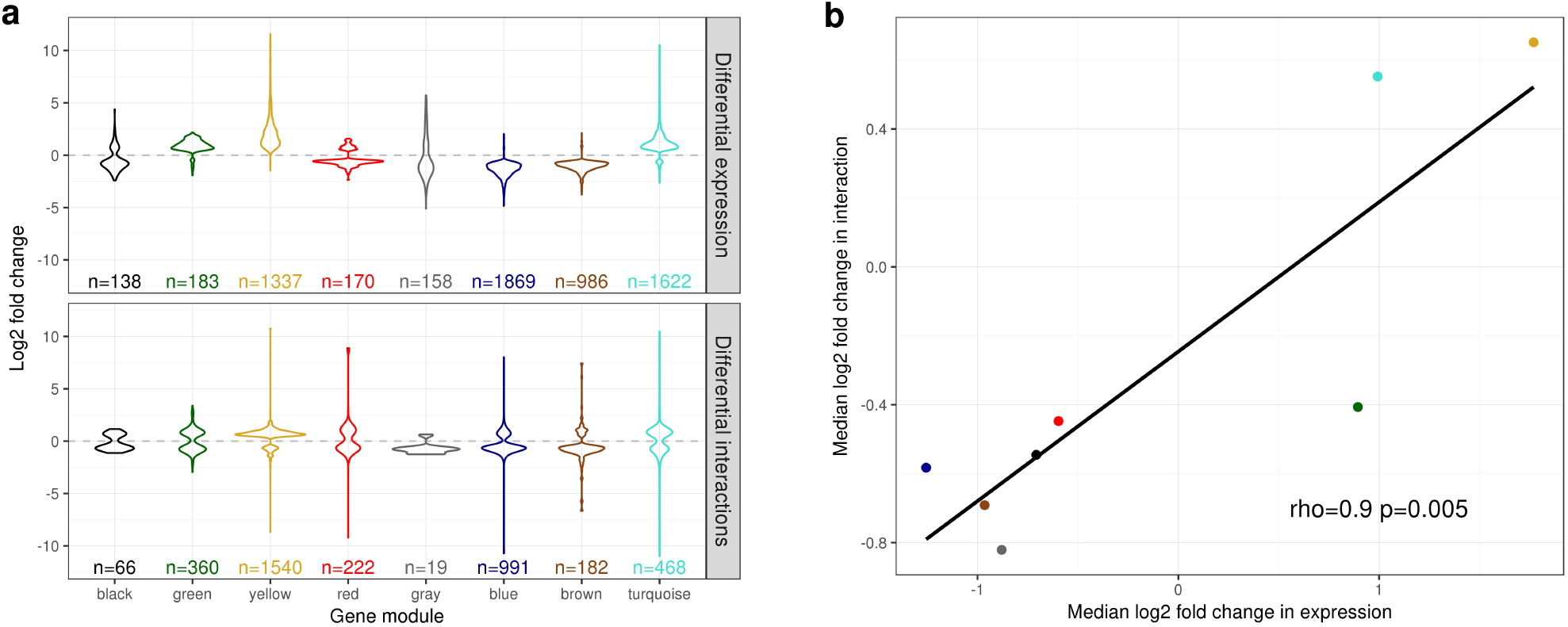
Change in PCHi-C interactions correlate with change in gene expression. **a** Distribution of significant (FDR<0.01) fold changes induced by activation of CD4^+^ T cells in (top) gene expression, and (bottom) differential PCHi-C interactions for differentially expressed genes in by module. **b** Median significant expression and interaction fold changes by module are correlated (Spearman rank correlation).

As we sequenced total RNA without a poly(A) selection step, we were able to detect regulatory region RNAs (regRNAs), which are generally non-polyadenylated and serve as a mark for promoter and enhancer activity[18]. We defined 6,147 “expressed” regRNAs (see Methods) that mapped within regulatory regions defined by a 15 state ChromHMM[19] model (Additional File 2: Fig. S6) and validated them by comparison to an existing cap analysis of gene expression (CAGE) dataset[20] which has been successfully used to catalog active enhancers.[21] We found 2,888/3,897 (74%) regRNAs expressed in non-activated cells overlap CAGE defined enhancers. This suggests that the combination of chromatin state annotation and total RNA-seq data presents an alternative approach to capture active enhancers.

Almost half (48%) of expressed regRNAs showed differential expression after activation (2,254/681 up/down regulated; FDR<0.01). To determine whether activity at these regRNAs could be related to that at PCHi-C linked genes, we focused attention on a subset of 640 intergenic regRNAs, which correspond to a definition of eRNAs[22]. Of these, 404 (63%) overlapped PIRs detected in CD4^+^ T cells and we found significant agreement in the direction of fold changes at eRNAs and their PCHi-C linked protein coding genes in activated CD4^+^ T cells (p<0.0001, Fig. 3b). We also observed a synergy between regRNA expression and the estimated effect of a PIR on expression with a gain or loss of a PIR overlapping a differentially regulated regRNA having the largest estimated effect on gene expression (Fig. 3c), supporting a sequential model of gene activation[23]. While regRNA function is still unknown[22], our results demonstrate the detection, by PCHi-C, of condition-specific connectivity between promoters and enhancers involved coordinating gene regulation.

**Fig. 3.**
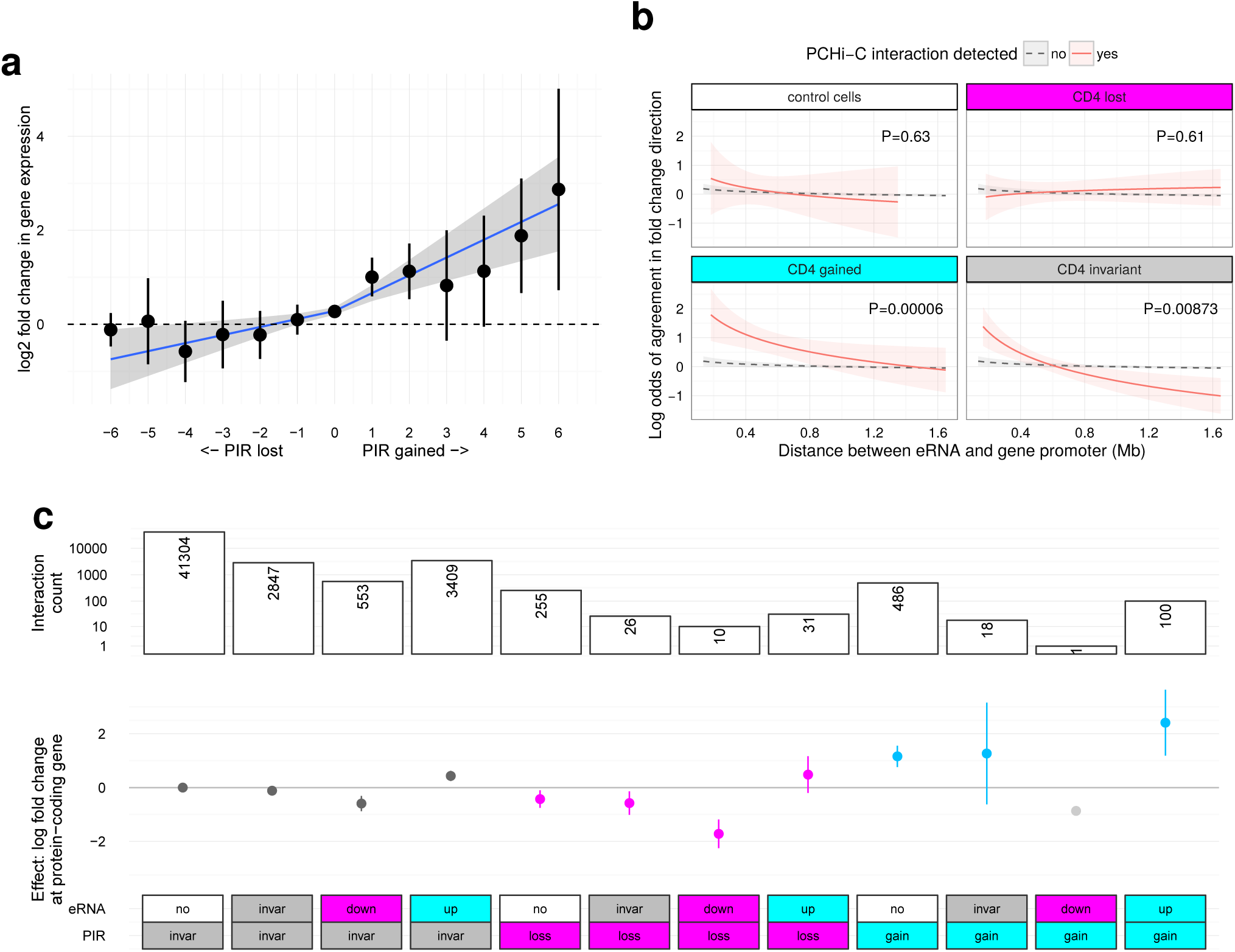
PCHi-C interactions and enhancer activity predict change in gene expression. **a** Change in gene expression at protein coding genes (log_2_ fold change, y axis) correlates with the number of PIRs gained or lost upon activation (x axis). **b** Fold change at transcribed sequence within the intergenic subset of regulatory regions (“eRNAs”) was more likely to agree with the direction of protein coding gene fold change when the two are linked by PCHi-C (red) in activated CD4^+^ T cells compared to pairs of eRNA and protein coding genes formed without regard to PCHi-C derived interactions (background, grey, p<10^-4^). Interactions were categorised as control (present only in megakaryocytes and erythroblasts, our control cells), invariant (“invar”; present in non-activated and activated CD4^+^ T cells), “loss” (present in non-activated but not activated CD4^+^ T cells, and significantly down-regulated at FDR<0.01) or “gain” (present in activated but not non-activated CD4^+^ T cells, and significantly up-regulated at FDR<0.01). **c** gain or loss of PIRs upon activation predicts change in gene expression, with the estimated effect more pronounced if accompanied by up- or down-regulation at an eRNA. Points show estimated effect on gene expression of each interaction and lines the 95% confidence interval. PIRs categorised as in b. eRNAs categorised as no (undetected), invariant (“invar”, detected in non-activated and activated CD4^+^ T cells, differential expression FDR>=0.01), up (up-regulated; FDR<0.01) or down (down-regulated; FDR<0.01). Bar plot (top) shows the number of interactions underlying each estimate. Note that eRNA=down, PIR=gain (light gray) has only one observation so no confidence interval can be formed and is shown for completeness only.

### PCHi-C-facilitated mapping of candidate disease causal genes

We defined an experimental framework to integrate PCHi-C interactions with GWAS data to map candidate disease causal genes for autoimmune diseases (**Fig. 4**). First, to confirm that PCHi-C interactions inform interpretation of autoimmune disease GWAS, we tested whether PIRs were enriched for autoimmune disease GWAS signals by testing for different distributions of GWAS p values in PIRs of activated or non-activated CD4^+^ T cells compared to non-lymphoid cells (megakaryocytes and erythroblasts) and then in PIRs of activated compared to non-activated CD4^+^ T cells. To perform the test, we used *blockshifter*, which accounts for correlation between (1) neighbouring variants in the GWAS data and (2) neighbouring *Hind*III fragments in the interacting data by rotating one dataset with respect to the other, as previously proposed [24]. This method appropriately controls type 1 error rates, in contrast to methods based on counting associated SNP/PIRs which ignore correlation, such as a Fisher’s exact test (Additional File 2: Fig. S7). We found autoimmune GWAS signals were enriched in CD4^+^ T cell PIRs compared to non-autoimmune GWAS signals (Wilcoxon p = 2.5x10^-7^) and preferentially so in activated *versus* non-activated cells (Wilcoxon p = 4.8x10^-5^; Fig. 4).

**Fig. 4.**
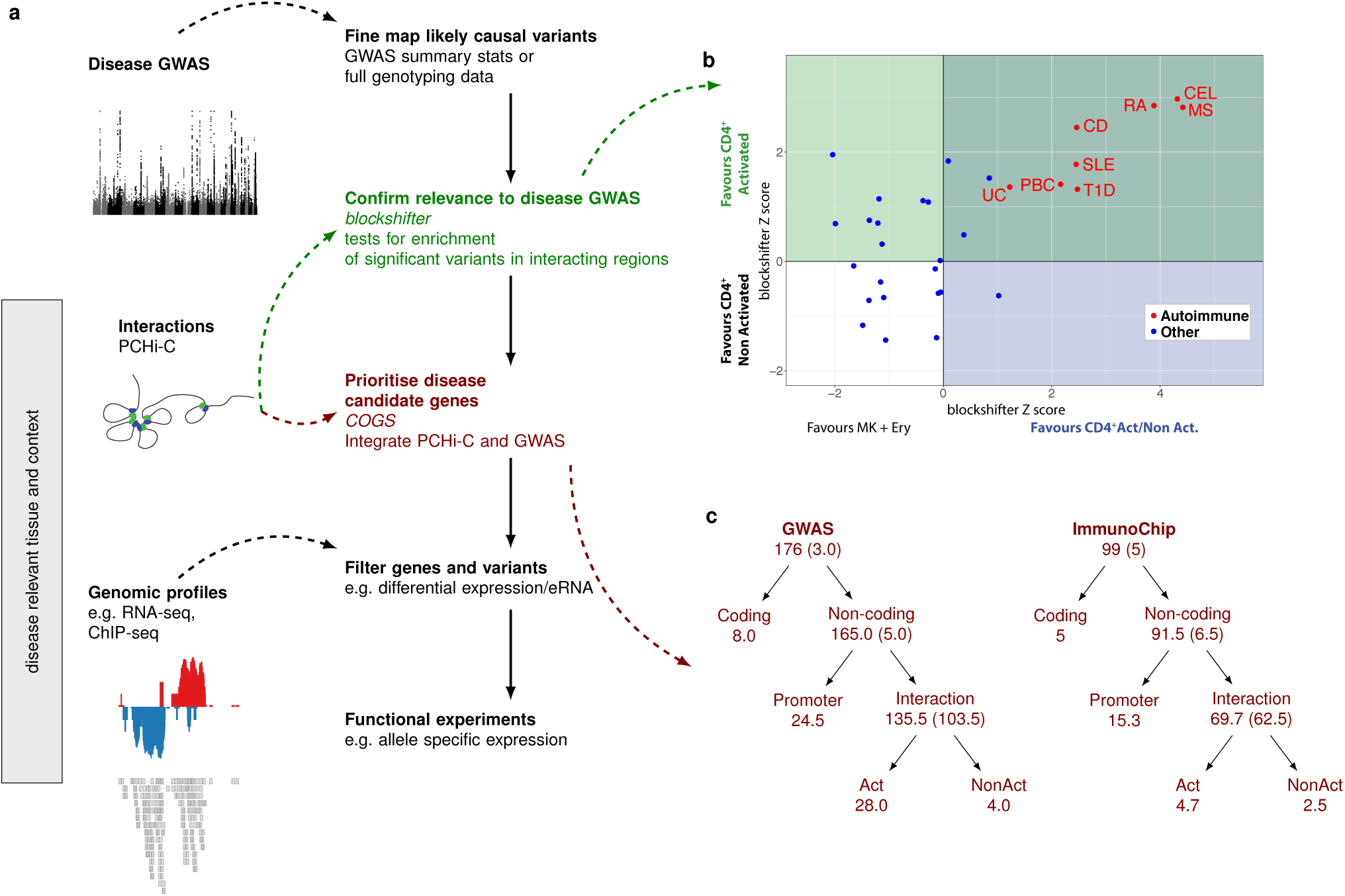
An experimental framework for identifying disease causal genes. **a** Before prioritising genes, enrichment of GWAS signals in PCHi-C interacting regions should be tested to confirm the tissue and context are relevant to disease. Then, probabilistic fine mapping of causal variants from the GWAS data can be integrated with the interaction data to prioritise candidate disease causal genes, a list which can be iteratively filtered using genomic datasets to focus on (differentially) expressed genes and variants which overlap regions of open or active chromatin. **b** Autoimmune disease GWAS signals are enriched in PIRs in CD4^+^ T cells generally compared to control cells (blockshifter Z score, x axis) and in PIRs in activated compared to non-activated CD4+ T cells (blockshifter Z score, y axis). Text labels correspond to datasets described in Additional File 6: Table S5. **c** Genes were prioritised with a COGS score>0.5 across five autoimmune diseases using genome-wide (GWAS) or targeted genotyping array (ImmunoChip) data. The numbers at each node give the number of genes prioritised at that level. Where there is evidence to split into one of two non-overlapping hypotheses (log_10_ ratio of gene scores>3), the genes cascade down the tree. Act and NonAct correspond to gene scores derived using PCHi-C data only in activated or non-activated cells, respectively. Where the evidence does not confidently predict which of the two possibilities is more likely, genes are “stuck” at the parent node (number given in brackets). When the same gene was prioritised for multiple diseases, we assigned fractional counts to each node, defined as the proportion of the n diseases for which the gene was prioritised at that node. Because of duplicate results between GWAS and ImmunoChip datasets, the total number of prioritised genes is 252 (see Table 1).

Next, we fine-mapped causal variants for five autoimmune diseases using genetic data from a dense targeted genotype array, the ImmunoChip (ATD, CEL, RA, T1D), and summary data from GWAS data imputed to 1000 Genomes Project (RA, SLE; Additional File 6: Table S5). For the ImmunoChip datasets, with full genotype data, we also imputed to 1000 Genomes and used a Bayesian fine mapping approach[25] which avoids the need for stepwise regression or assumptions of single causal variants and which provides a measure of posterior belief that any given variant is causal by aggregating posterior support over models containing that variant. Variant-level results are given in Additional Files 7-8: Tables S6a-S6b, and show that of 73 regions with genetic association signals to at least one disease (minimum p < 5x10^-8^; 106 disease associations), nine regions have strong evidence that they contain more than one causal variant (posterior probability > 0.5), among them the well studied region on chromosome 10 containing the candidate gene *IL2RA[25]*. For the GWAS summary data, we make the simplifying assumption that there exists a single causal variant in any LD-defined genetic region and again generate posterior probabilities that each variant is causal[26]. To integrate these variant level data with PCHi-C interactions and prioritize protein coding genes as candidate causal genes for each autoimmune disease, we calculated gene-level posterior support by summing posterior probabilities over all models containing variants in PIRs for a given gene promoter, within the promoter fragment or within its immediate neighbour fragments. Neighbouring fragments are included because of limitations in the ability of PCHi-C to detect very proximal interactions (within a region consisting of the promoter baited fragment and one *Hind*III fragment either side). The majority of gene scores were close to 0 (99% of genes have a score <0.05) and we chose to use a threshold of 0.5 to call genes prioritised for further investigation. Having both ImmunoChip and summary GWAS data for RA allowed us to compare the two methods. Overlap was encouraging: of 24 genes prioritised for RA from ImmunoChip, 20 had a gene score > 0.5 in the GWAS prioritisation, a further three had gene scores > 0.2. The remaining gene, *MDN1,* corresponded to a region which has a stronger association signal in the RA-ImmunoChip than RA-GWAS dataset, which may reflect the greater power of direct genotyping versus imputation, given that the RA-ImmunoChip signal is mirrored in ATD and T1D (Additional File 2: Fig. S8). We prioritised a total of 245 unique protein coding genes, 108 of which related to activation sensitive interactions (Additional Files 9-10: Tables S7a-7b, Fig. 4). Of 118 prioritised genes which could be related through interactions to a known susceptibility region, 63 (48%) lay outside that disease susceptibility region. The median distance from peak signal to prioritised gene was 153 kb. Note that prioritisation can be one (variant)-to-many (genes) because a single PIR can interact with more than one promoter, and promoter fragments can contain more than one gene promoter. Note also that the score reflects both PCHi-C interactions and the strength and shape of association signals (Additional File 2: Fig. S9), therefore a subset of prioritised genes relate to an aggregation over sub-genomewide significant GWAS signals. This is therefore a “long” list of prioritised genes which requires further filtering (**Table 1**). One hundred and seventy nine (of 245) prioritised genes were expressed in at least one activation state; we highlight specifically the subset of 118 expressed genes which can be related to a genome-wide significant GWAS signal through proximity of a genome-wide significant SNP (p<5x10^-8^) to a PIR. Of these, 82 were differentially expressed, 48 related to activation-sensitive interactions and 63 showed overlap of GWAS fine-mapped variants with an expressed eRNA (Additional File 9: Table S7a).

**Table 1.** Number of genes prioritised for autoimmune disease susceptibility under successive filters.

| Group | Description | Number of genes |
| --- | --- | --- |
| 1 | Total | 245 |
| 2 | ... Expressed | 179 |
| 3 | ... Proximal GWAS significant SNP ($p < 5 \times 10^{-8}$) | 118 |
| 4 | ... Prioritised gene differentially expressed upon activation | 82 |
| 5 | ... Prioritisation relates to activation sensitive interactions | 48 |
| 6 | ... GWAS signal overlaps expressed regRNA in at least one state | 63 |

Note that group 2 is a subset of group 1, group 3 is a subset of group 2, and groups 4, 5 and 6 are all subsets of group 3 but not necessarily of each other.

Taken together, our results reflect the complexity underlying gene regulation, and the context-driven effects that common autoimmune disease-associated variants may have on candidate genes. Our findings are consistent with, and extend, previous observations[7, 8] and we highlight six examples which exemplify both activation-specific and activation-invariant interactions.

### Exemplar regions

Here we highlight specific examples of prioritised genes with plausible autoimmune disease candidacy which illustrate three characteristics we found frequently, namely (1) the identification of candidate genes some distance from association signals, and (2) the tendency for multiple gene promoters to be identified as interacting with the same sets of disease associated variants and (3) genes prioritised in only one state of activation.

As an example of the first, CEL has been associated with a region on chromosome 1q31.2, for which *RGS1* has been named as a causal candidate due to proximity of associated variants to its promoter[27]. Sub-genome-wide significant signals for T1D (min. p=1.5x10^-6^) across the same SNPs which are associated with CEL have been interpreted as a colocalising T1D signal in the region[28]. *RGS1* has recently been shown to have a role in the function of T follicular helper cells in mice[29], the frequencies of which and their associated IL-21 production have been shown to be increased in T1D patients[30]. However, our analysis also prioritises, in activated T cells, the strong functional candidate genes *TROVE2* and *UCHL5*, over half a megabase distant and with three intervening genes not prioritised (**Fig. 5**). *UCHL5* encodes ubiquitin carboxyl-terminal hydrolase-L5 a deubiquitinating enzyme that stabilizes several Smad proteins and TGFBR1, key components of the TGF-beta1 signalling pathway[31, 32]. *TROVE2* is significantly upregulated upon activation (FDR=0.005) and encodes Ro60, an RNA binding protein that indirectly regulates type-I IFN-responses by controlling endogenous Alu RNA levels[33]. A global anti-inflammatory effect for *TROVE2* expression would fit with its effects on gut (CEL) and pancreatic islets (T1D).

**Fig. 5.**
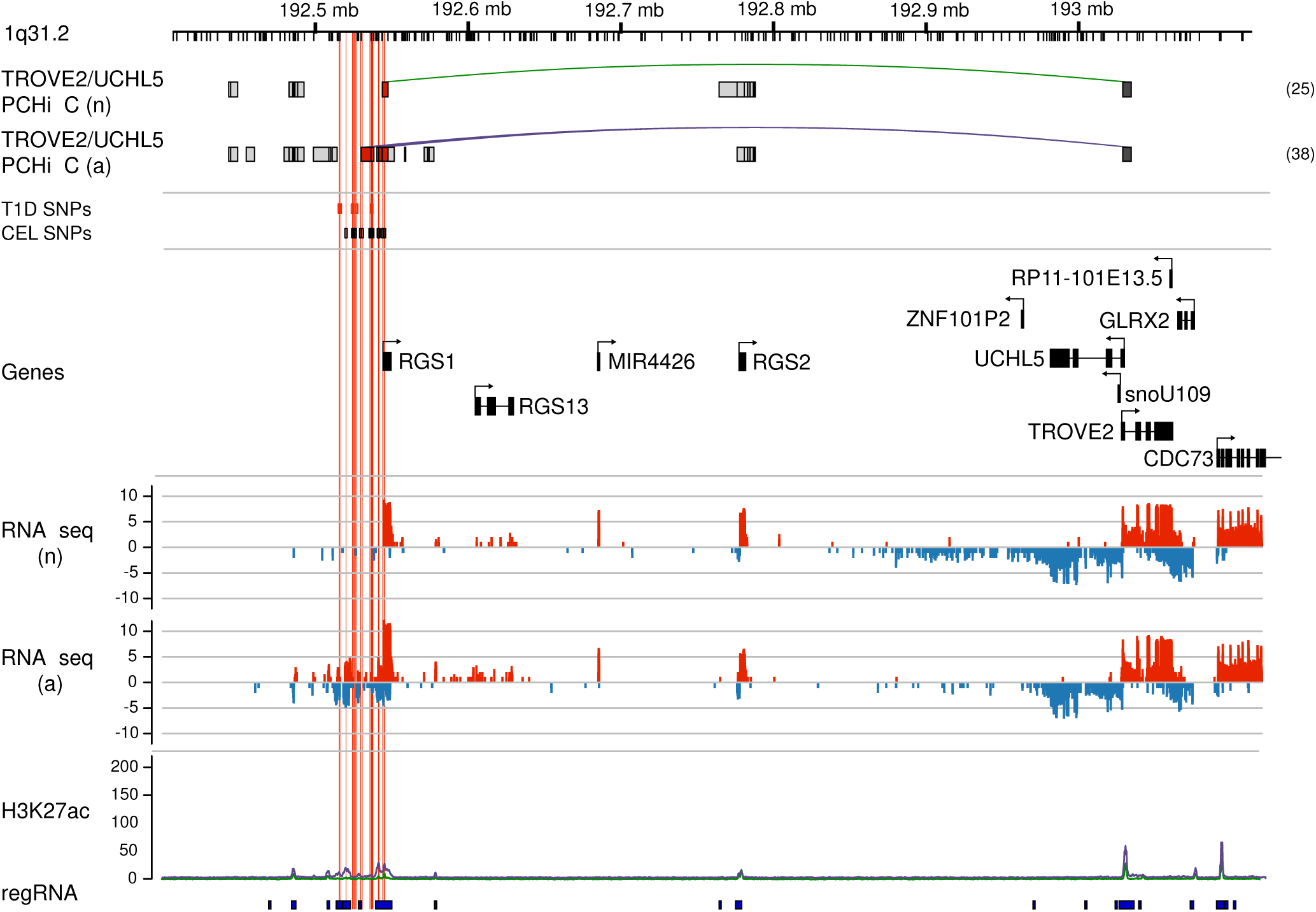
*TROVE2* and *UCLH5* on chromosome 1 are prioritised for CEL. The ruler shows chromosome location, with HindIII sites marked by ticks. The top tracks show PIRs for prioritised genes in non-activated (n) and activated (a) CD4^+^ T cells. Green and purple lines are used to highlight those PIRs containing credible SNPs from our fine mapping. The total number of interacting fragments per PCHi-C bait is indicated in parentheses for each gene in each activation state. Dark grey boxes indicate promoter fragments; light grey boxes, PIRs containing no disease associated variants; and red boxes, PIRs overlapping fine mapped disease associated variants. The position of fine mapped variants area indicated by red boxes and vertical red lines. Gene positions and orientation (ensembl v75) are shown above log2 read counts for RNA-seq forward (red) and reverse (blue) strand. H3K27ac background-adjusted read count is shown in non-activated (green line) and activated (purple line) and boxes on the regRNA track show regions considered through ChromHMM to have regulatory marks.

A similar situation is seen in the chromosome 1q32.1 region associated with T1D in which *IL10* has been named as a causal candidate gene[34]. Together with *IL10*, prioritised through proximity of credible SNPs to the *IL10* promoter, we prioritised other *IL10* gene family members *IL19*, *IL20* and *IL24* as well as two members of a conserved three-gene immunoglobulin-receptor cluster (*FCMR* and *PIGR,* Additional File 2: Fig. S10). While *IL19, IL20* and *PIGR* were not expressed in CD4^+^ T cells, *FCMR* was down- and *IL24* and *IL10* were up-regulated following CD4^+^ T cell activation. IL-10 is recognised as an important anti-inflammatory cytokine in health and disease[35] and candidate genes *FCMR* and *IL24* are components of a recently identified proinflammatory module in Th17 cells[36].

This, region also exemplifies characteristic 2: candidate causal variants lay in *Hind*III fragments that interacted with multiple genes. Parallel results have demonstrated co-regulation of multiple PCHi-C interacting genes by a single variant[37], suggesting that disease related variants may act on multiple genes simultaneously, consistent with the finding that regulatory elements can interact with multiple promoters[38–40]. This region also shows that clusters of multiple adjacent PIRs can be detected for the same promoter. It remains to be further validated whether all PIRs detected within such clusters correspond to ‘causal’ interactions or whether some such PIRs are ‘bystanders’ of strong interaction signals occurring in their vicinity. The use of PCHi-C nonetheless adds considerable resolution compared to simply considering topologically associating domains (TADs), which have a median length of 415 kb in naive CD4^+^ T cells[37] compared to a median of 37.5 kb total PIR length per gene in non-activated CD4^+^ T cells (Additional File 2: Fig. S11).

Multiple neighbouring genes were also prioritised on chromosome 16q24.1: *EMC8*, *COX4I1* and *IRF8*, the last only in activated T cells, for two diseases: RA and SLE (Additional File 2: Fig. S12). Candidate causal variants for SLE and RA fine-mapped to distinct PIRs, yet all these PIRs interact with the same gene promoters, suggesting that interactions, possibly specific to different CD4^+^ T cell subsets, may allow us to unite discordant GWAS signals for related diseases[6, 41, 42]. *EMC8* and *COX4I1* RNA expression was relatively unchanged by activation, whereas *IRF8* expression was upregulated 97-fold, coincident with the induction of 16 intergenic *IRF8* PIRs, four of which overlap autoimmune disease fine-mapped variants. Although the dominant effect of *IRF8* is to control the maturation and function of the mononuclear phagocytic system[43], a T cell-intrinsic function regulating CD4^+^ Th17 differentiation has been proposed[44]. Our data further link the control of Th17 responses to susceptibility to autoimmune disease including RA and SLE[45].

Another notable example *AHR,* was one of the 38 genes we prioritised in only one state of activation (characteristic 3, **Fig. 4**): for rheumatoid arthritis (RA), *AHR* was prioritised specifically in activated T cells rather than non-activated T cells (Additional File 2: Fig. S13). *AHR* is a high affinity receptor for toxins in cigarette smoke that has been linked to RA previously through differential expression in synovial fluid of patients, though not through GWAS[46]. Our analysis prioritises *AHR* for RA due to a sub-genome-wide significant signal (rs71540792, p=2.9x10^-7^) and invites attempts to validate the genetic association in additional RA patients.

### Interaction-mediated regulation of *IL2RA* expression

Given our prior interest in the potential for autoimmune-disease associated genetic variants to regulate *IL2RA* expression [42], we were interested to see PCHi-C interactions between some of these variants and the *IL2RA* promoter, and attempted to confirm the predicted functional effects on *IL2RA* expression experimentally. *IL2RA* encodes CD25, a component of the key trimeric cytokine receptor that is essential for high-affinity binding of IL-2, regulatory T cell survival and T effector cell differentiation and function[47]. Multiple variants in and near *IL2RA* have been associated with a number of autoimmune diseases[34, 48–50]. We have previously fine-mapped genetic causal variants for T1D and multiple sclerosis (MS) in the *IL2RA* region[25], identifying five groups of SNPs in intron 1 and upstream of *IL2RA,* each of which is likely to contain a single disease causal variant. Out of the group of eight SNPs previously denoted “A”[25], three (rs12722508, rs7909519 and rs61839660) are located in an area of active chromatin in intron 1, within a PIR that interacts with the *IL2RA* promoter in both activated and non-activated CD4^+^ T cells (Fig. 6a). These three SNPs are also in LD with rs12722495 (r^2^>0.86) that has previously been associated with differential surface expression of CD25 on memory T cells[42] and differential responses to IL-2 in activated Tregs and memory T cells[51]. We measured the relative expression of *IL2RA* mRNA in five individuals heterozygous across all group “A” SNPs and homozygous across most other associated SNP groups, in a four-hour activation time-course of CD4^+^ T cells. Allelic imbalance was observed consistently for two reporter SNPs in intron 1 and in the 3’ UTR in non-activated CD4^+^ T cells in each individual (Fig. 6b; Additional File 2: Fig. S14a), validating a functional effect of the PCHi-C-derived interaction between this PIR and the *IL2RA* promoter in non-activated CD4^+^ T cells. While the allelic imbalance was maintained in non-activated cells cultured for 2-4 hours, the imbalance was lost in cells activated under our *in vitro* conditions. Since increased CD25 expression with rare alleles at group “A” SNPs has previously been observed on memory CD4+ T cells but not the naive or Treg subsets that are also present in the total CD4+ T cell population[42], we purified memory cells from 8 group “A” heterozygous individuals and confirmed activation-induced loss of allelic imbalance of IL2RA mRNA expression in this more homogeneous population (Fig. 6c, Additional File 2: Fig. S14b; Wilcoxon p=0.007). *IL2RA* is one of the most strongly upregulated genes upon CD4+ T cell activation, showing a 65-fold change in expression in our RNA-seq data. Concordant with the genome-wide pattern (Fig. 3), the *IL2RA* promoter fragment gains PIRs that accumulate H3K27ac modifications upon activation and these, as well as potentially other changes marked by an increase in H3K27ac modification at rs61839660 and across the group A SNPs in intron 1, could account for the loss of allelic imbalance. These results emphasise the importance of steady-state CD25 levels on CD4^+^ T cells for the disease association mediated by the group A SNPs, levels which will make the different subsets of CD4^+^ T cells more or less sensitive to the differentiation and activation events caused by IL-2 exposure *in vivo[52]*.

**Fig. 6.**
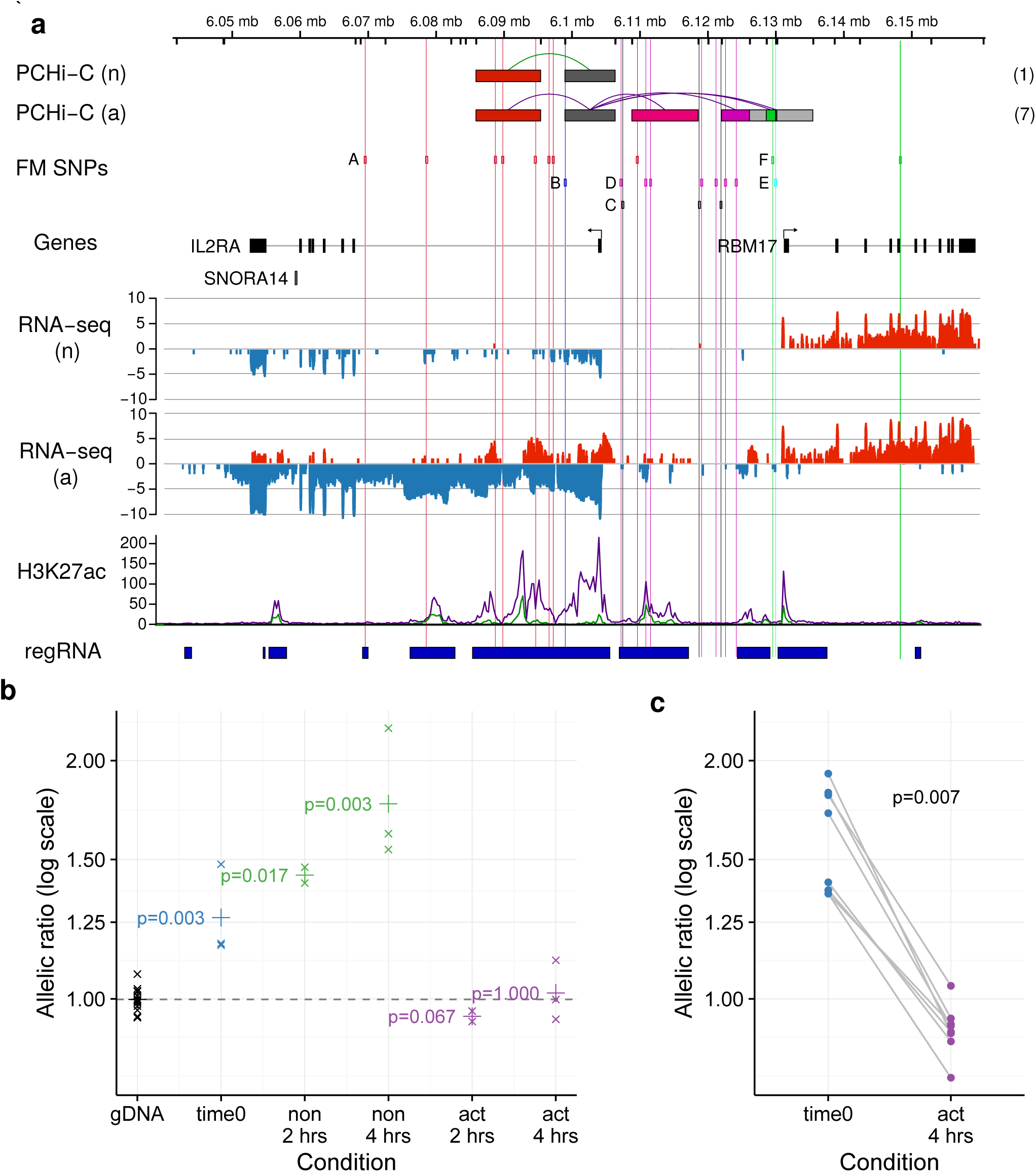
PCHi-C interactions link the *IL2RA* promoter to autoimmune disease associated genetic variation, which leads to expression differences in *IL2RA* mRNA. **a** The ruler shows chromosome location, with HindIII sites marked by ticks. The top tracks show PIRs for prioritised genes in non-activated (n) and activated (a) CD4^+^ T cells. Green and purple lines are used to highlight those PIRs containing credible SNPs for the autoimmune diseases T1D and MS fine mapped on chromosome 10p15. Six groups of SNPs (A-F) highlighted in Wallace et al.[25] are shown, although note that group B was found unlikely to be causal. The total number of interacting fragments per PCHi-C bait is indicated in parentheses for each gene in each activation state. Dark grey boxes indicate promoter fragments; light grey boxes, PIRs containing no disease associated variants; and coloured boxes, PIRs overlapping fine mapped disease associated variants. PCHi-C interactions link a region overlapping group A in non-activated and activated CD4+ T cells to the *IL2RA* promoter (dark grey box) and regions overlapping groups D and F in activated CD4+ T cells only. RNA-seq reads (log_2_ scale, red=forward strand, blue=reverse strand) highlight the upregulation of IL2RA expression upon activation and concomitant increases in H3K27ac (non-activated, n, green line; activated, a, purple line) in the regions linked to the *IL2RA* promoter. Red vertical lines mark the positions of the group A SNPs. Numbers in parentheses show the total number of IL2RA PIRs detected in each state. Here we show those PIRs proximal to the *IL2RA* promoter. Comprehensive interaction data can be viewed at \url{http://www.chicp.org. **b** Allelic imbalance in mRNA expression in total CD4+ T cells from individuals heterozygous for group A SNPs using rs12722495 as a reporter SNP in non-activated (non) and activated (act) CD4^+^ T cells cultured for 2 or 4 hours, compared to genomic DNA (gDNA, expected ratio=1). Allelic ratio is defined as the ratio of counts of T to C alleles. ‘x’=geometric mean of the allelic ratio over 2-3 replicates within each of 4-5 individuals, and p values from a Wilcoxon rank sum test comparing cDNA to gDNA are shown. `+’ shows the geometric mean allelic ratio over all individuals. **c** Allelic imbalance in mRNA expression in memory CD4^+^ T cells differs between ex vivo (time 0) and four hour activated samples from eight individuals heterozygous for group A SNPs using rs12722495 as a reporter SNP. p value from a paired Wilcoxon signed rank test is shown.

### Discussion

Our results illustrate the changes in chromosome conformation detected by PCHi-C in a single cell type in response to a single activation condition. That the PCHi-C technique can indeed link enhancers to their target genes is supported by our evidence that the direction of fold changes at eRNAs is connected to those at their PCHi-C linked protein-coding genes. Our results also provide support for the candidacy of certain genes and sequences in GWAS regions as causal for disease. Our results illustrate the changes in chromosome conformation in a single cell type in response to a single activation condition, in addition to providing support for the candidacy of certain genes and sequences in GWAS regions as causal for disease. That the PCHi-C technique can indeed link enhancers to their target genes is supported by our evidence that the direction of fold changes at eRNAs is connected to those at their PCHi-C linked protein coding genes. Recent attempts to link GWAS signals to variation in gene expression in primary human cells have sometimes found only limited overlap[53–55]. One explanation may be that these experiments miss effects in specific cell subsets or states, especially given the transcriptional diversity between the many subsets of memory CD4^+^ T cells[56]. We highlight the complex nature of disease association at the *IL2RA* region where additional PIRs for *IL2RA* gained upon activation overlap other fine-mapped disease-causal variants (Fig. 6a), suggesting that other allelically-imbalanced states may exist in activated cells, which may also correspond to altered disease risk. For example, the PIR containing rs61839660, a group A SNP, also contains an activation eQTL for *IL2RA* expression in CD4^+^ T effectors[57] marked by rs12251836, which is unlinked to the group A variants and was not associated with T1D[57]. Furthermore, rs61839660 itself has recently been reported as a QTL for methylation of the *IL2RA* promoter as well as an eQTL for *IL2RA* expression in whole blood[55, 58]. The differences between CD25 expression in different T cell subsets[59, 60], and the rapid activation-induced changes in gene and regulatory expression, chromatin marks and chromosome interactions we observe, imply that a large diversity of cell types and states will need to be assayed to fully understand the identity and effects of autoimmune disease causal variants.

Our approach, like others such as eQTL analyses and integration of GWAS variants with chromatin state information, offers a view of disease through the prism of purified cell subsets in specific states of activation. However, a more complete range of cell types and activation states will be needed for the comprehensive understanding of complex diseases for which multiple cell types are aetiologically involved. It will be challenging to assay this greater diversity of cell types and states in the large numbers of individuals needed for traditional eQTL studies, particularly for cell-type or condition-specific eQTLs that have been shown to generally have weaker effects[61, 62]. Allele-specific expression (ASE) is a more powerful design to quantify the effects of genetic variation on gene expression with modest sample sizes[63] and the targeted ASE approach that we adopt enables testing of individual variants or haplotypes at which donors are selected to be heterozygous, while controlling for other potentially related variants at which donors are selected to be homozygous.

### Conclusions

Here we have presented an approach for connecting disease-associated variants derived from GWAS with putative target genes based on promoter interactome maps obtained with PCHi-C. By using statistical fine mapping of GWAS data, integrated with PCHi-C, to highlight both likely disease causal variants and their potential target genes, we enable the design of targeted ASE analyses for functional confirmation of individual effects. This systematic experimental framework offers an alternative approach to candidate causal gene identification for variants with cell state-specific functional effects, with achievable sample sizes.

## Methods

### CD4^+^ T cell purification and activation, preparation for genomics assays

Blood samples were obtained from healthy donors selected from the Cambridge BioResource. Donors were excluded if they were diagnosed with autoimmune disease or cancer, were receiving immunosuppressants, cytotoxic agents or intravenous immunoglobulin or had been vaccinated or received antibiotics in the 2-4 weeks preceding the blood donation. CD4^+^ T cells were isolated from whole blood using RosetteSep (STEMCELL technologies, Canada) according to the manufacturer’s instructions. Purified CD4^+^ T cells (average =96.5% pure, range 92.9 - 98.7%) were washed in X-VIVO 15 supplemented with 1% AB serum (Lonza, Switzerland) and penicillin/streptomycin (Invitrogen, UK) and plated in 96-well CELLSTAR U-bottomed plates (Greiner Bio-One, Austria) at a concentration of 2.5 x 10^5^ cells / well. Cells were left untreated or stimulated with Dynabeads human T activator CD3/CD28 beads (Invitrogen, UK) at a ratio of 1 bead: 3 cells for 4 hours at 37°C and 5% CO. Cells were harvested, centrifuged, supernatant removed and either, (i) resuspended in RLT buffer (RNeasy micro kit, Qiagen, Germany) for RNA-seq (0.75-1 x 10^6^ CD4^+^ T cells / pool and activation state) (ii) fixed in formaldehyde for capture Hi-C (44-101 x 10^6^ CD4^+^ T cells / pool and activation state) or ChIP-seq (16-26 x 10^6^ CD4^+^ T cells / pool and activation state) as detailed in[37].

ChIP-seq (H3K27ac, H3K4me1, H3K4me3, H3K27me3, H3K9me3, H3K36me3) was carried out according to BLUEPRINT protocols[64]. Formaldehyde fixed cells were lysed, sheared and DNA sonicated using a Bioruptor Pico (Diagenode). Sonicated DNA was pre-cleared (Dynabeads Protein A, Thermo Fisher) and ChIP performed using BLUEPRINT validated antibodies and the IP-Star automated platform (Diagenode). Libraries were prepared and indexed using the iDeal library preparation kit (Diagenode) and sequenced (Illumina HiSeq, paired-end).

For PCHi-C[37], DNA was digested overnight with *Hind*III, end labeled with biotin-14-dATP and ligated in preserved nuclei. De-crosslinked DNA was sheared to an average size of 400 bp, end-repaired and adenine-tailed. Following size selection (250-550 bp fragments), biotinylated ligation fragments were immobilized, ligated to paired-end adaptors and libraries amplified (7-8 PCR amplification rounds). Biotinylated 120-mer RNA baits targeting both ends of *Hind*III restriction fragments that overlap Ensembl-annotated promoters of protein-coding, noncoding, antisense, snRNA, miRNA and snoRNA transcripts were used to capture targets[65]. After enrichment, the library was further amplified (4 PCR cycles) and sequenced on the Illumina HiSeq 2500 platform. Each PCHi-C library was sequenced over 3 lanes generating 50 bp paired-end reads.

### PCHi-C interaction calls

Raw sequencing reads were processed using the HiCUP pipeline[66] and interaction confidence scores were computed using the CHiCAGO pipeline[14] as previously described[37]. We considered the set of interactions with high confidence scores (> 5) in this paper.

Raw PCHi-C read counts from 3 replicates and 2 conditions were transformed into a matrix, and a trimmed mean of M-values normalization was applied to account for library size differences. Subsequently, a voom normalization was applied to log-transformed counts in order to estimate precision weights per contact, and differential interaction estimates were obtained after fitting a linear model on a paired design, using the limma Bioconductor R package[67].

### Microarray measurement of gene expression

We recruited 20 healthy volunteers from the Cambridge BioResource. Total CD4^+^ T cells were isolated from whole blood within 2 hours of venepuncture by RosetteSep (StemCell technologies). To assess the transcriptional variation in response to TCR stimulation, 10^6^ CD4^+^ T cells were cultured in U-bottom 96-well plates in the presence or absence of human T activator CD3/CD28 beads at a ratio of 1 bead: 3 cells. Cells were harvested at 2, 4, 6 or 21 hours post-stimulation, or after 0, 6 or 21 hours in the absence of stimulation. Three samples from the 6 hour unstimulated time point were omitted from the study due to insufficient cell numbers, and a further four samples were dropped after quality control, resulting in a total of 133 samples that were included in the final analysis. RNA was isolated using the RNAeasy kit (Qiagen) according to the manufacturer’s instructions.

cDNA libraries were synthesized from 200 ng total RNA using a whole-transcript expression kit (Ambion) according to the manufacturer’s instructions and hybridized to Human Gene 1.1 ST arrays (Affymetrix). Microarray data were normalized using a variance stabilizing transformation[68] and differential expression was analysed in a paired design using limma[67]. Genes were clustered into modules using WGCNA[69].

### ChIP sequencing and regulatory annotation

ChIP sequencing reads for all histone modification assays and control experiments were mapped to the reference genome using BWA-MEM[70], a Burrows-Wheeler genome aligner. Samtools[71] was employed to filter secondary and low-quality alignments (we retained all read pair alignments with PHRED score > 40 that matched all bits in SAM octal flag 3, and did not match any bits in SAM octal flag 3840). The remaining alignments were sorted, indexed and a whole-genome pileup was produced for each histone modification, sample and condition triple.

We used ChromHMM[19], a multivariate hidden Markov model, to perform a whole-genome segmentation of chromatin states for each activation condition. First, we binarized read pileups for each chromatin mark pileup using the corresponding control experiment as a background model. Second, we estimated the parameters of a 15-state hidden Markov model (a larger state model resulted in redundant states) using chromosome 1 data from both conditions. Parameter learning was re-run five times using different random seeds to assess convergence. Third, a whole-genome segmentation was produced for each condition by running the obtained model on the remaining chromosomes. Each state from the obtained model was manually annotated, and states indicating the presence of promoter or enhancer chromatin tags were selected (E4-E11, Additional File 2: Fig. S6). Overlapping promoter or enhancer regions in non-activated and activated genome segmentations were merged to create a CD4^+^ T cell regulatory annotation. Thus, we defined 53,534 regulatory regions (Additional Files 11-13: Tables S8a-S8c).

### RNA sequencing

Total RNA was isolated using the RNeasy kit (Qiagen) and the concentrations and integrity were quantified using Bioanalyzer (Agilent); all samples reached RINs > 9.8. Two pools of RNA were generated from three and four donors and for each experimental condition. cDNA libraries were prepared from 1ug total RNA using the stranded NEBNext Ultra Directional RNA kit (New England Biolabs), and sequenced on HiSeq (Illumina) at an average coverage of 38 million paired-end reads/pool. RNA sequencing reads were trimmed to remove traces of library adapters by matching each read with a library of contaminants using Cutadapt[72], a semi-global alignment algorithm. Owing to our interest in detecting functional enhancers, which constitute transcription units on their own right, we mapped reads to the human genome using STAR[73], a splicing-aware aligner. This frees us from relying on a transcriptome annotation which would require exact boundaries and strand information for all features of interest, something not available in case of promoters and enhancers.

After alignment, we employed Samtools[71] to discard reads with an unmapped pair, secondary alignments and low-quality alignments. The resulting read dataset, with an average of 33 million paired-end reads/sample, was sorted and indexed. We used FastQC (v0.11.3, http://www.bioinformatics.babraham.ac.uk/projects/fastqc/) to ensure all samples had regular GC content (sum of deviations from normal includes less than 15% of reads), base content per position (difference A vs T and G vs C less than 10% at all positions) and kmer counts (no imbalance of kmers with p < 0.01) as defined by the tool. We augmented Ensembl 75 gene annotations with regulatory region definitions obtained from our ChIP-seq analysis described above, and defined them as present in both genome strands due to their bidirectional transcription potential. For each RNA-seq sample, we quantified expression of genomic and regulatory features in a two-step strand-aware process using HTSeq[74]. For each gene we counted the number of reads that fell exactly within its exonic regions, and did not map to other genomic elements. For each regulatory feature we counted the number of reads that fell exactly within its defined boundaries, and did not map to other genomic or regulatory elements.

By construction, this quantification scheme counts each read at most once towards at most one feature. Furthermore, strand information is essential to be able to assign reads to features in regions with overlapping annotations. For example, distinguishing intronic eRNAs from pre-mRNA requires reads originating from regulatory activity in the opposite strand from the gene.

Feature counts were transformed into a matrix, and a trimmed mean of M-values normalization was applied to account for library size differences, plus a filter to discard features below an expression threshold of < 0.4 counts per million mapped reads in at least two samples, a rather low cutoff, to allow for regulatory RNAs to enter differential expression calculations. This threshold equates to approximately 15 reads, given our mapped library sizes of ∼35 million paired-end reads. A voom normalization was applied to log-transformed counts in order to estimate precision weights per gene, and differential expression estimates were obtained after fitting a linear model on a paired design, using the limma Bioconductor R package[67]. There was a strong correlation (rho=0.81) between microarray and RNA-seq fold change estimates at 4 hours.

### Comparison of regRNAs to FANTOM CAGE data

We compared expressed regRNA regions detected in our non-activated CD4^+^ T cell samples versus those found using CAGE-seq by the FANTOM5 Consortium. RNA-seq, using a regulatory reference obtained from chromatin states, yields 17,175 features expressed with at least 0.4 counts per million in both non-activated CD4^+^ T cell samples. Among those, 3,897 correspond to regulatory regions. Unstimulated CD4^+^ samples from FANTOM5 (<u>http://fantom.gsc.riken.jp/5/datafiles/latest/basic/human.primary_cell.hCAGE/</u>, samples 10853, 11955 and 11998) contain 266,710 loci expressed (with at least one read) in all 3 samples.

We found 13,178 of our 17,175 expressed CD4^+^ T cell features overlap expressed loci in CAGE data (77%). Conversely, 243,596/266,710 CAGE loci overlap CD4^+^ T cell features (91%). Similarly 2,888/3,897 expressed regRNAs overlap expressed loci in CAGE data (74%).

### Comparison of PCHi-C and ChIA-PET interactions

We downloaded supplementary table 1 from <u>http://www.nature.com/cr/journal/v22/n3/extref/cr201215x1.xlsx [15]</u> and counted the overlaps of PCHi-C interactions from CD4^+^ T cells and comparitor cells (megakaryoctyes and erythroblasts) in distance bins. R code to replicate the analysis is at <u>https://github.com/chr1swallace/cd4-pchic/blob/master/chepelev.R</u>. Calling interactions requires correction for the expected higher density of random collisions at shorter distances[75] which are explicitly modelled by CHICAGO[14] used in this study but not in the ChIA-PET study[15]. As a result, we expected a higher false positive rate from the ChIA-PET data at shorter distances.

### Regression of gene expression against PIR count and eRNA expression

We related measures of gene expression (absolute log2 counts or log2 fold change) to numbers of PIRs or numbers of PIRs overlapping specific features using linear regression. We used logistic regression to relate agreement between fold change direction at PCHi-C linked protein coding genes and eRNAs. We used robust clustered variance estimates to account for the shared baits for some interactions across genes with the same prey. Enrichment of chromatin marks in interacting baits and prey were assessed by logistic regression modelling of a binary outcome variable (fragment overlapped specific chromatin peak) against a fragment width and a categorical explanatory variable (whether the *Hind*III fragment was a bait or prey and the cell state the interaction was identified in), using block bootstrapping of baited fragments (<u>https://github.com/chr1swallace/genomic.autocorr</u>) to account for spatial correlation between neighbouring fragments.

### GWAS summary statistics

We used a compendium of 31 GWAS datasets [37] (Additional File 6: Table S5). Briefly we downloaded publicly available GWAS statistics for 31 traits. Where necessary we used the *liftOver* utility to convert these to GRCh37 coordinates. To remove spurious association signals, we removed variants with P< 5 x 10^-8^ for which there were no variants in LD (r^2^>0.6 using 1000 genomes EUR cohort as a reference genotype panel) or within 50 kb with P<10^-5^. We masked the MHC region (GRCh37:chr6:25-35Mb) from all downstream analysis due to its extended LD and known strong and complex associations with autoimmune diseases.

Comparison of GWAS data and PIRs requires dense genotyping coverage. For GWAS which did not include summary statistics imputed for non-genotyped SNPs, we used a poor man’s imputation (PMI) method [37] to impute. We imputed p values at ungenotyped variants from 1000 Genomes EUR phase 3 by replacing missing values with those of their nearest proxy variant with r^2^>0.6, if one existed. Variants that were included in the study but did not map to the reference genotype set were also discarded.

To calculate posterior probabilities that each SNP is causal under a single causal variant assumption, we divided the genome into linkage disequilibrium blocks of 1cM based on data from the HapMap project (<u>http://hapmap.ncbi.nlm.nih.gov/downloads/recombination/2011-01_phaseII_B37/</u>). For each region excluding the MHC we used code modified from *Giambartolomei et al.[76]* to compute approximate Bayes factors for each variant using the Wakefield approximation[77], and thus posterior probabilities that each variant was causal as previously proposed[78]. The method assumes a normal prior on the population log relative risk centred at 0, and we set the variance of this distribution to 0.04, equivalent to a 95% belief that the true relative risk is between 0.66 and 1.5 at any causal variant. We set the prior probability that any variant is causal for disease to 10^-4^.

### Testing of the enrichment of GWAS summary statistics in PIRs using *blockshifter*

We used the *blockshifter* method [37] (<u>https://github.com/ollyburren/CHIGP</u>) to test for a difference between variant posterior probability distributions in *Hind*III fragments with interactions identified in test and control cell types using the mean posterior probability as a measure of central location. *Blockshifter* controls for correlation within the GWAS data due to LD and interaction restriction fragment block structure by employing a rotating label technique similar to that described in GoShifter[24] to generate an empirical distribution of the difference in means under the null hypothesis of equal means in the test and control set. Runs of one or more PIRs (separated by at most one *Hind*III fragment) are combined into ‘blocks’, that are labeled unmixed (either test or control PIRs) or mixed (block contains both test and control PIRs). Unmixed blocks are permuted in a standard fashion by reassigning either test or control labels randomly, taking into account the number of blocks in the observed sets. Mixed blocks are permuted by conceptually circularising each block and rotating the labels. A key parameter is the gap size - the number of non-interacting *Hind*III fragments allowed within a single block, with larger gaps allowing for more extended correlation.

We used simulation to characterise the type 1 error and power of *blockshifter* under different conditions and to select an optimal gap size. Firstly, from the Javierre *et al.* dataset[37] *we selected a test (Activated or Non Activated CD4+ T Cells) and control (Megakaryocyte or Erythroblast) set of PIRs with a CHiCAGO score > 5, as a reference set for blockshifter* input.

Using the European 1000 genomes reference panel, we simulated GWAS summary statistics, under different scenarios of GWAS/PIR enrichment. We split chromosome 1 into 1cM LD blocks and used reference genotypes to compute a covariance matrix for variants with minor allele frequency above 1%, Σ. GWAS Z scores can be simulated as multivariate normal with mean μ and variance Σ[79]. Each block may contain no causal variants (GWAS_null_, μ = 0) or one (GWAS_alt_). For GWAS_alt_ blocks, we pick a single causal variant, *i*, and calculate the expected non-centrality parameter (NCP) for a 1 degree of freedom chi-square test of association at this variant and its neighbours. This framework is natural because the NCP at any variant *j* can be expressed as the NCP at the causal variant multiplied by the r^2^ between variants *i* and *j[80]*. In each case we set the NCP at the causal variant to 80 to ensure that each causal variant was genome-wide significant (P < 5 x 10^-8^). μ is defined as the square root of this constructed NCP vector.

For all scenarios we randomly chose 50 GWAS_alt_ blocks leaving the remaining 219 GWAS_null_. Enrichment is determined by the preferential location of simulated causal variants within test PIRs. In all scenarios, each causal variant has a 50% chance of lying within a PIR, to mirror a real GWAS in which we expect only a proportion of causal variants to be regulatory in any given cell type. Under the enrichment-null scenario, used to confirm control of type 1 error rate, the remaining variants were assigned to PIRs without regard for whether they were identified in test or control tissues. To examine power, we considered two different scenarios with PIR-localised causal variants chosen to be located specifically in test PIRs with either 50% probability, scenario power (1), or 100%, scenario power (2). Note that a PIR from the test set may also be in the control set, thus, as with a real GWAS, not all causal variants will be informative for this test of enrichment.

For each scenario we further considered variable levels of genotyping density, corresponding to full genotyping (everything in 1000 Genomes), HapMap imputation (the subset of SNPs also in Stahl et al. [81] dataset) or genotyping array (the subset of SNPs also on the Illumina 550k array). Where genotyping density is less than full, we used our proposed poor man’s imputation (PMI) strategy to fill in Z scores for missing SNPs.

We ran *blockshifter*, with 1000 null permutations, for each scenario and PMI condition for 4000 simulated GWAS, with a *blockshifter* superblock gap size parameter (the number of contiguous non-PIR *Hind*III fragments allowed within one superblock) of between 1 and 20 and supplying numbers of cases and controls from the RA dataset[49].

For comparison we also investigated the behaviour of a naive test for enrichment for the null scenario. We computed a 2x2 table variants according to test and control PIR overlap, and whether a variant’s posterior probability of causality exceeded an arbitrary threshold of 0.01, and Fisher’s exact test to test for enrichment.

### Enrichment of GWAS summary statistics in CD4+ and activated CD4+ PIRs

We compared the following sets using all GWAS summary statistics, with a superblock gap size of 5 (obtained from simulations above) and 10,000 permutations under the null:-

- Total CD4^+^ Activated + Total CD4^+^ NonActivated (test) versus Endothelial precursors + Megakaryocytes (control)
- Total CD4^+^ Activated (test) versus Total CD4^=^ NonActivated (control).

### Variant posterior probabilities of inclusion, full genotype data (ImmunoChip)

We carried out formal imputation to 1000 Genomes Project EUR data using IMPUTE2 [82] and fine-mapped causal variants in each of the 179 regions where a minimum p < 0.0001 was observed using a stochastic search method which allows for multiple causal variants in a region, (https://github.com/chr1swallace/GUESSFM)[25]. Despite the pre-selection of regions associating with autoimmune diseases on the ImmunoChip, we chose to again set the prior probability that any variant was causal to 10^-4^, to align our analysis with that applied to the GWAS summary data. The prior probability for individual models follows a binomial distribution, according to the number of causal variants represented, so that the prior for each of the 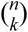 *k* SNP causal variant models was (10^-4^)^*k*^ (1-10^-4^)^(*n-k*)^ where *n* is the number of SNPs considered in the region. The posterior probabilities for models that contained variants which overlapped PIRs for each gene were aggregated to compute PIR-level marginal posterior probabilities of inclusion.

### Variant posterior probabilities of inclusion, summary statistics

Where we have only summary statistics of GWAS data already imputed to 1000 Genomes, we divided the genome into linkage disequilibrium blocks of 0.1cM based on data from the HapMap project (<u>http://hapmap.ncbi.nlm.nih.gov/downloads/recombination/2011-01_phaseII_B37/</u>). For each region excluding the MHC we use code modified from *Giambartolomei et al.[76]* to compute approximate Bayes factors for each variant using the Wakefield approximation[77], and thus posterior probabilities that each variant was causal assuming at most one causal variant per region as previously proposed[78].

### Computation of gene prioritisation scores

We used the COGS method[37] (<u>https://github.com/ollyburren/CHIGP</u>) to prioritise genes for further analysis. We assign variants to the first of the following three categories it overlaps for each annotated gene, if any

1. coding variant: the variant overlaps the location of a coding variant for the target gene.
2. promoter variant: the variant lies in a region baited for the target gene or adjacent restriction fragment.
3. PIR variant: the variant lies in a region overlapping any PIR interacting with the target gene.

We produced combined gene/category scores by aggregating, within LD blocks, over models with a variant in a given set of PIRs (interacting regions), or over *Hind*III fragments baited for the gene promoter and immediate neighbours (promoter regions), or over coding variants to generate marginal probabilities of inclusion (MPPI) for each hypothesised group. We combine these probabilities across LD blocks, *i*, using standard rules of probability to approximate the posterior probability that at least one LD block contains a causal variant:

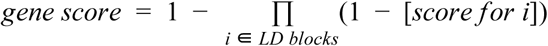

Thus the score takes a value between 0 and 1, with 1 indicating the strongest support. We report all results with score > 0.01 in Additional File 9: Table S7b, but focus in this manuscript on the subset with scores > 0.5.

Because COGS aggregates over multiple signals, a gene may be prioritised because of many weak signals or few strong signals in interacting regions. To predict the expected information for future users of this method, we considered the subset of 76 input regions with genome-wide significant signals (p<5x10^-8^) in ImmunoChip datasets. We prioritised at least one gene with a COGS score > 0.5 in 35 regions, with a median of three genes/region (interquartile range, IQR = 1.5-4). Equivalent analysis of the genome-wide significant GWAS signals prioritised a median of two genes/region (interquartile range = 1-3). This suggests that this algorithm might be expected to prioritise at least one gene in about half the genomewide significant regions input when run on a relevant cell type.

Whilst components 1 and 2 are fixed for a given gene and trait the contribution of variants overlapping PIRs varies depending on the tissue context being examined. We developed a hierarchical heuristic method to ascertain for each target gene which was the mostly likely component and cell state. Firstly for each gene we compute the gene score due to genic variants (components 1 + 2) and variants in PIRs (component 3) using all available tissue interactions for that gene. We use the ratio of the genic score to PIRs score in a similar manner to a Bayes factor to decide whether causal variants contributing to the gene score are more likely to lie within the gene or within its associated PIRs. If a genic location is more likely (gene.score ratio >3) we iterate and compare if the gene score due to coding variants (component 1) is more likely than for promoter variants (component 2). Similarly if PIRs are more likely we compare PIR gene scores for activated vs non-activated cells. If at any stage no branch is substantially preferred over its competitor (ratio of gene scores < 3) we return the previous set as most likely, otherwise we continue until a single cell state/set is chosen. In this way we can prioritize genes based on the overall score and label as to a likely mechanism for candidate causal variants.

### Allele-specific expression assays

Total CD4^+^ T cells were isolated from five healthy donors and activated as described above and were harvested after 0, 2 and 4 hours in RLT Plus buffer. Selected donors were heterozygous at all eight group A SNP and, homozygous for group C and F SNPs. Two and three of the donors were homozygous for the group D and E SNP groups, respectively (Additional File 14: Table S9). Memory CD4^+^ T cells were sorted from cryopreserved PBMC from an additional eight healthy donors as viable, αβ TCR^+^, CD4^+^, CD45RA^-^,CD127^+^, CD27^+^ cells using a FACSAria III cell sorter (BD Biosciences). Sorted cells were either activated for 4 hours in culture as described above or resuspended directly in RLT plus buffer post-sort. Total RNA was extracted using Qiagen RNeasy Micro plus kit and cDNA was synthesised using Superscript III reverse transcriptase (Thermo Fisher) according to manufacturer’s instructions. To perform allele-expression experiments we used a modified version of a previously described method for quantifying methylation in bisulfite sequence data[83]. A two-stage PCR was used, the first round primers were designed to flank the variant of interest using Primer3 (http://bioinfo.ut.ee/primer3-0.4.0/primer3/) and adaptor sequences were added to the primers (Sigma), shown as lowercase letters (rs61839660_ASE_F tgtaaaacgacggccagtGCACACACCTATCCTAGCCT, rs61839660_ASE_R caggaaacagctatgaccCCCACAGAATCACCCACTCT, product size 114bp; rs12244380_ASE_F tgtaaaacgacggccagtTTCGTGGGAGTTGAGAGTGG, rs12244380_ASE_R caggaaacagctatgaccTTAAAAGAGTTCGCTGGGCC, product size 180bp; rs12722495_ASE_F tgtaaaacgacggccagtGTGAGTTTCAATCCTAAGTGCGA, rs12722495_ASE_R caggaaacagctatgaccATTAAGCGGACTCTCTGGGG, product size 97bp). The first round PCR contains 10 μl of Qiagen multiplex PCR mastermix, 0.5 μl of 10 nmol forward primer, 0.5 μl of 10 nmol reverse primer, 4 μl of cDNA and made up to 20 μl with ultra-pure water. The PCR cycling conditions were 95°C for 15 minutes hot start, followed by 30 cycles of the following steps: 95°C for 30 seconds, 60°C for 90 seconds and 72°C for 60 seconds, finishing with a 72°C for 10 minutes cycle. The first round PCR product was cleaned using AmpureXP beads (Beckman Coulter) according to manufacturer’s instructions. To add Illumina sequence compatible ends to the individual first round PCR amplicons, additional primers were designed to incorporate P1 and A sequences plus sample-specific index sequences in the A primer, through hybridisation to adapter sequence present on the first round gene-specific primers. Index sequences are as published[83]. The second-round PCR contained 8 μl of Qiagen multiplex PCR mastermix, 2.0 μl of ultra-pure water, 0.35 μl of each forward and reverse index primer, 5.3 μl of Ampure XP-cleaned first-round PCR product. The PCR cycling conditions were 95°C for 15 minutes hotstart, followed by 7 cycles of the following steps: 95°C for 30 seconds, 56°C for 90 seconds, 72°C for 60 seconds, finishing with 72°C for 10 minutes cycle. All PCR products were pooled at equimolar concentrations based on quantification on the Shimadzu Multina. AmpureXP beads were used to remove unincorporated primers from the product pool. We used the Kapa Bioscience library quantification kit to accurately quantify the library according to manufacturer’s instructions before sequencing on an Illumina MiSeq v3 reagents (2 x 300 bp reads).

### Statistical analysis of allele-specific expression data

Sequence data was processed using the Methpup package (<u>ttps://github.com/ollyburren/Methpup</u>) to extract counts of each allele at rs12722495, and rs12244380 (Additional File 15: Table S10). Individuals were part of a larger cohort genotyped on the ImmunoChip and were phased using snphap (<u>https://github.com/chr1swallace/snphap</u>) to confirm which allele at each SNP was carried on the same chromosome as A2=rs12722495:C or A1=rs12722495:T. Allelic imbalance was quantified as the ratio A2/A1 and was averaged across replicates within individuals using a geometric mean. Allelic ratios in cDNA and gDNA were compared using Wilcoxon rank sum tests. P values are shown in Fig. 6b and Additional File 2: Fig. S14. Full details are in <u>https://github.com/chr1swallace/cd4-pchic/blob/master/IL2RA-ASE.R</u>.

## Declarations

### Ethics approval and consent to participate

All samples and information were collected with written and signed informed consent. The study was approved by the local Peterborough and Fenland research ethics committee for the project entitled: ‘An investigation into genes and mechanisms based on genotype-phenotype correlations in type 1 diabetes and related diseases using peripheral blood mononuclear cells from volunteers that are part of the Cambridge BioResource project’ (05/Q0106/20). Experimental methods comply with the Helsinki Declaration.

### Consent for publication

Not applicable

### Availability of data and materials

The datasets generated and/or analysed during the current study are available in repositories or additional files as indicated below:

- PCHi-C data are available as raw sequencing reads from EGA (<u>https://www.ebi.ac.uk/ega)</u> accession number EGAS00001001911 or CHICAGO-called interactions (Additional File 5: Table S4) and are available for interactive exploration via <u>http://www.chicp.org</u>
- RNA-seq and ChIPseq data are available as raw sequencing reads from EGA (<u>https://www.ebi.ac.uk/ega</u>) accession number EGAS00001001961.
- Microarray data are available at ArrayExpress, <u>https://www.ebi.ac.uk/arrayexpress</u>, accession number E-MTAB-4832
- Processed datasets are available as Additional Files
- Code used to analyse the data are available from <u>https://github.com/chr1swallace/cd4-pchic</u> except where other URLs are indicated in Methods. The version of scripts used in the production of this manuscript are archived under <u>https://doi.org/10.5281/zenodo.832849</u>.

### Author Contributions

Study conceived by: CW, JAT, LSW, PF, and led by CW. Interpreted the data: CW, OSB, AJC, ARG, DR, LSW, JAT. Sequence data analysis: ARG. HiCUP analysis: SW. ASE experiments and analysis: DR. Microarray experiments and analysis: XCD,RCF, RC, CW. Statistical analysis: OSB, JC, NJC, CW, ARG. Laboratory experiments: AJC, BJ, DR, JJL, FB, SPR, KD. Wrote the paper: CW, OSB, AJC, ARG and contributed to writing: JAT, LSW, MS. Revised the paper: all authors. Genetic association data processing: CW, OSB, ES. Supervised capture Hi-C experiments: MS and PF. Supervised cell experiments: AJC, MF, WO, PF, JAT and LW.

## Acknowledgements

We thank all study participants and family members.

We thank the Wellcome Trust for funding the AITD UK national collection; all doctors and nurses in Birmingham, Bournemouth, Cambridge, Cardiff, Exeter, Leeds, Newcastle and Sheffield for recruitment of patients and J. Franklyn, S. Pearce (Newcastle) and P. Newby (Birmingham) for preparing and providing DNA samples on Graves’ disease patients.

This research utilizes resources provided by the Type 1 Diabetes Genetics Consortium, a collaborative clinical study sponsored by the National Institute of Diabetes and Digestive and Kidney Diseases (NIDDK), National Institute of Allergy and Infectious Diseases (NIAID), National Human Genome Research Institute (NHGRI), National Institute of Child Health and Human Development (NICHD), and JDRF and supported by U01 DK062418.

We gratefully acknowledge the participation of all Cambridge NIHR BioResource volunteers, and thank the Cambridge BioResource staff for their help with volunteer recruitment. We thank the National Institute for Health Research Cambridge Biomedical Research Centre and NHS Blood and Transplant for funding. Further information can be found at www.cambridgebioresource.org.uk

We thank the High-Throughput Genomics Group at the Wellcome Trust Centre for Human Genetics (funded by Wellcome Trust grant reference 090532/Z/09/Z) for the generation of the sequencing data.

We thank Stephen Eyre for helpful comments on the manuscript, and N. Soranzo and the HaemGen consortium for sharing blood trait GWAS summary statistics.

The authors acknowledge the assistance and support of the National Institute for Health Research (NIHR) Cambridge Biomedical Research Centre. Helen Stevens, Meeta Maisuria-Armer, Pamela Clarke, Gillian Coleman, Sarah Dawson, Simon Duley, Jennifer Denesha and Trupti Mistry for sample processing. Judy Brown, Lynne Adshead, Amie Ashley, Anna Simpson and Niall Taylor for laboratory administration and procurement support. Vin Everett and Sundeep Nanuwa for logistical and web development.

We thank investigators of published ImmunoChip studies for making available their raw genotyping data (David van Heel, celiac disease; Stephen Eyre, rheumatoid arthritis; Matthew Simmonds, Stephen Gough, Jayne Franklyn, and Oliver Brand, autoimmune thyroid disease).

## Additional Files

- Additional File 1: additional-file-1-tableS1-gene-modules.csv.gz (CVS) Table S1 Gene Modules inferred from WGCNA analysis of microarray timecourse. Expression fold changes and associated false discovery rates (adjusted p values) are from RNA-seq data at the 4 hour timepoint.
- Additional File 2: additional-file-2-figures-s1-s15.pdf (PDF) Figures S1-S15
- Additional File 3: additional-file-3-tableS2-rnaseq.csv.gz (CSV) Table S2 Results of differential expression analysis on RNA-seq data. Features are defined in the GTF file in Additional File 11: Table S8a.
- Additional File 4: additional-file-4-tableS3-HindIII-baits-e75.bed.gz (BED) Table S3 Baited HindIII fragments used for capture of Hi-C libraries, annotated with Ensembl annotated genes.
- Additional File 5: additional-file-5-tableS4-interactions.csv.gz (CSV) Table S4 PCHi-C interactions called with the CHiCAGO pipeline. Annotation for baited fragments is given in Additional File 4: Table S3. PIRs are called other ends (“oe”). CHICAGO scores for activated (“Total_CD4_Activated”) and non-activated (“Total_CD4_NonActivated”) CD4+ T cells were considered called with confidence if above 5. We also conducted differential analysis, and the read counts input into that are given by the columns P1.non - P3.act, with the results summarised by their log fold change (logFC) and FDR. Bait-PIR pairs are shown only if the CHiCAGO score is >=5 for at least one CD4+ T cell.
- Additional File 6: additional-file-6-tableS5-gwas-ic-meta-data.xslx (XLSX) Table S5 Summary of GWAS data used. ‘type’ indicates whether the trait was quantitative (QUANT) or case/control (CC). For CC, ‘cases’ and ‘controls’ columns represent the number of individuals included in the study, while for QUANT, the number of individuals is given in the cases column. ‘Category’ indicates broader classes of traits.
- Additional File 7: additional-file-7-tableS6a-fine-mapping-ichip.csv.gz (CSV) Table S6a Results of ImmunoChip fine mapping by GUESSFM
- Additional File 8: additional-file-8-tableS6b-fine-mapping-gwas.csv.gz (CSV) Table S6b Results of GWAS summary statistic fine mapping
- Additional File 9: additional-file-9-tableS7a-COGS-05.csv.gz (CSV) Table S7a Autoimmune disease COGS gene prioritisation. Overall COGS gene scores (COGS_Overall_Gene_Score) for each gene and autoimmune disease are shown together with the prioritised category and score associated with that category (COGS_Category_Gene_Score) (Fig. 3). The ‘analysis’ column describes whether the input data was GWAS or ImmunoChip (ICHIP) and whether summary statistic (SS) or GUESSFM (GF) fine mapping was used. ‘diff.expr’ indicates whether the gene was not expressed (NA) or, if expressed, whether there was differential expression at the FDR<0.01 level (up, down, or nsig). Similarly, ‘diff.erna’ indicates whether the HindIII fragment containing the strongest SNP signal is differentially expressed with the same categories. Using data from ImmunoBase (http://www.immunobase.org - accessed 06/06/2016) we annotate genes near (within 5Mb) previously reported disease susceptibility regions, with contextual annotation ‘Closest_Disease_Susceptibility_Region’, ‘Closest_Min_P_Value_Susceptibility_SNP’, ‘Closest_Min_P_Value_Susceptibility_SNP_P_Value’ ‘PIR_Overlaps_Disease_DSR’ indicates that the PIR driving the prioritisation for a gene/disease overlaps an ImmunoBase known disease susceptibility region for that trait. Restricted to the subset of genes with scores > 0.5 that are analysed in this paper.
- Additional File 10: additional-file-10-tableS7b-COGS-complete.csv.gz (CSV) Table S7b As above, complete results.
- Additional File 11: additional-file-11-tableS8a-genfeatures.gtf.gz (CSV) Table S8a GTF file with definitions for all Ensembl 75 genomic features plus CD4-specific regulatory regions inferred from chromatin states. These regulatory regions have been named with identifiers containing a CD4R prefix, assigned a regulatory biotype, and marked as pertaining to both genomic strands due to their bidirectional transcription potential.
- Additional File 12: additional-file-12-tableS8b-segmentation-act.bed.gz (BED) Table S8b whole-genome segmentation of non-activated and activated CD4 T cells into 15 states obtained from a CHROMHMM analysis using ChIP-seq data for activated CD4^+^ T cells
- Additional File 13: additional-file-13-tableS8c-segmentation-non.bed.gz (BED) Table S8c whole-genome segmentation of non-activated and activated CD4 T cells into 15 states obtained from a CHROMHMM analysis using ChIP-seq data for non-activated CD4^+^ T cells
- Additional File 14: additional-file-14-tableS9-IL2RA-donor-genotypes.xlsx (XLSX) Table S9. Genotypes for donors in the IL2RA ASE experiment across SNP groups A, C, D, E, F.
- Additional File 15: additional-file-15-tableS10-IL2RA-ASE.csv.gz (CSV) Table S10. Read counts for each allele at the IL2RA ASE experiment. The column Expt denotes sample id; time, the timepoint (0, 120, 240 minutes); stim, the condition (genomic DNA, time0 cDNA, stimulated or unstimulated cells cDNA.

## Supplementary Material

**Fig. S1:**
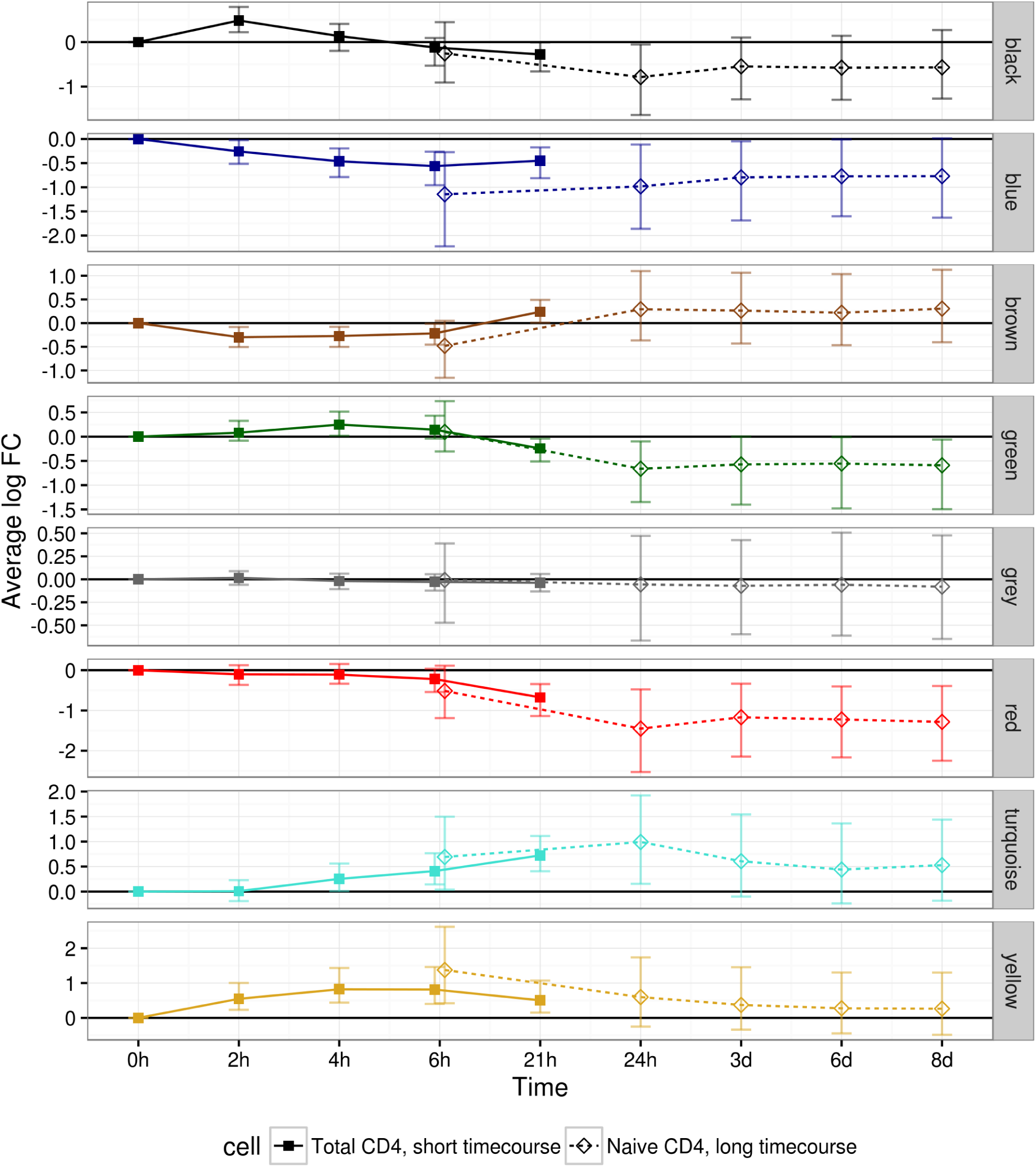
Comparison of longer and shorter CD4^+^ T cell activation timecourses. Microarray timecourse summary from this experiment (solid points) overplotted with a longer timecourse from GSE60680 (open points, Gustafsson et al, 2015). Points show the median log fold change amongst genes assigned to each module at each timepoint, with the interquartile range displayed as vertical ranges around each point. The results at 6 hours are slightly horizontally offset to allow the results from the two experiments to be visually distinguished. Note the non-linear mapping of time to the x axis, which contains a mixture of hours (h) and days (d), to allow visualization of the early timepoints in particular.

**Fig. S2:**
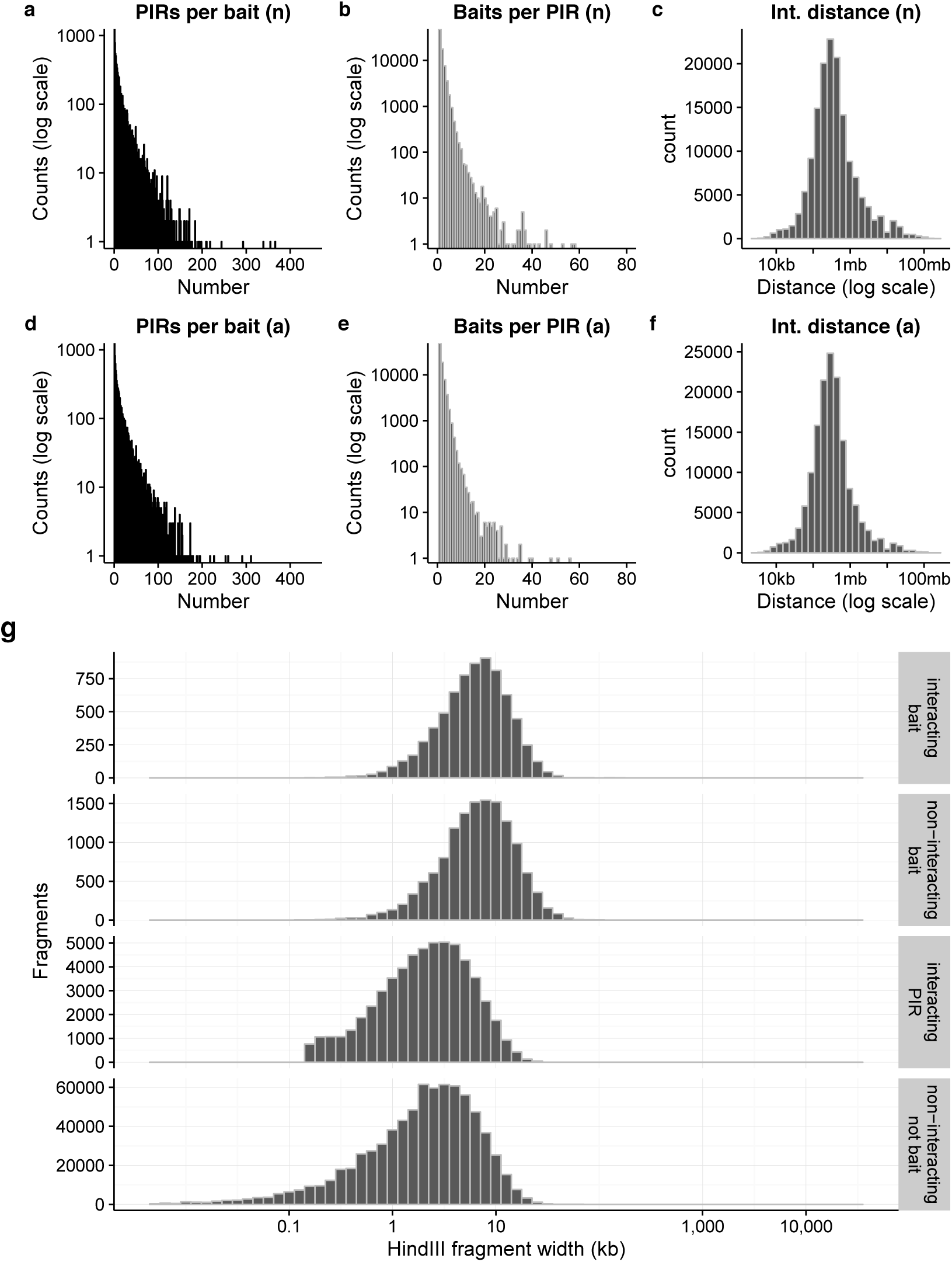
Summary distributions of interacting fragments. Distributions of **a**, **d** number of interacting promoter bait fragments per PIR; **b**, **e** PIRs per promoter fragment; and **c**, **f** distance between midpoints of promoter and PIR *Hind*III fragments in activated (**a-c**) and non-activated (**d-f**) CD4^+^ T cells. **f** Width profile of *Hind*III fragments according to whether they were baited promoter fragments or not, and interacting fragments or not. **g** *Hind*III fragment length in the four categories of interacting and non-interacting baited fragments and PIRs.

**Fig. S3:**
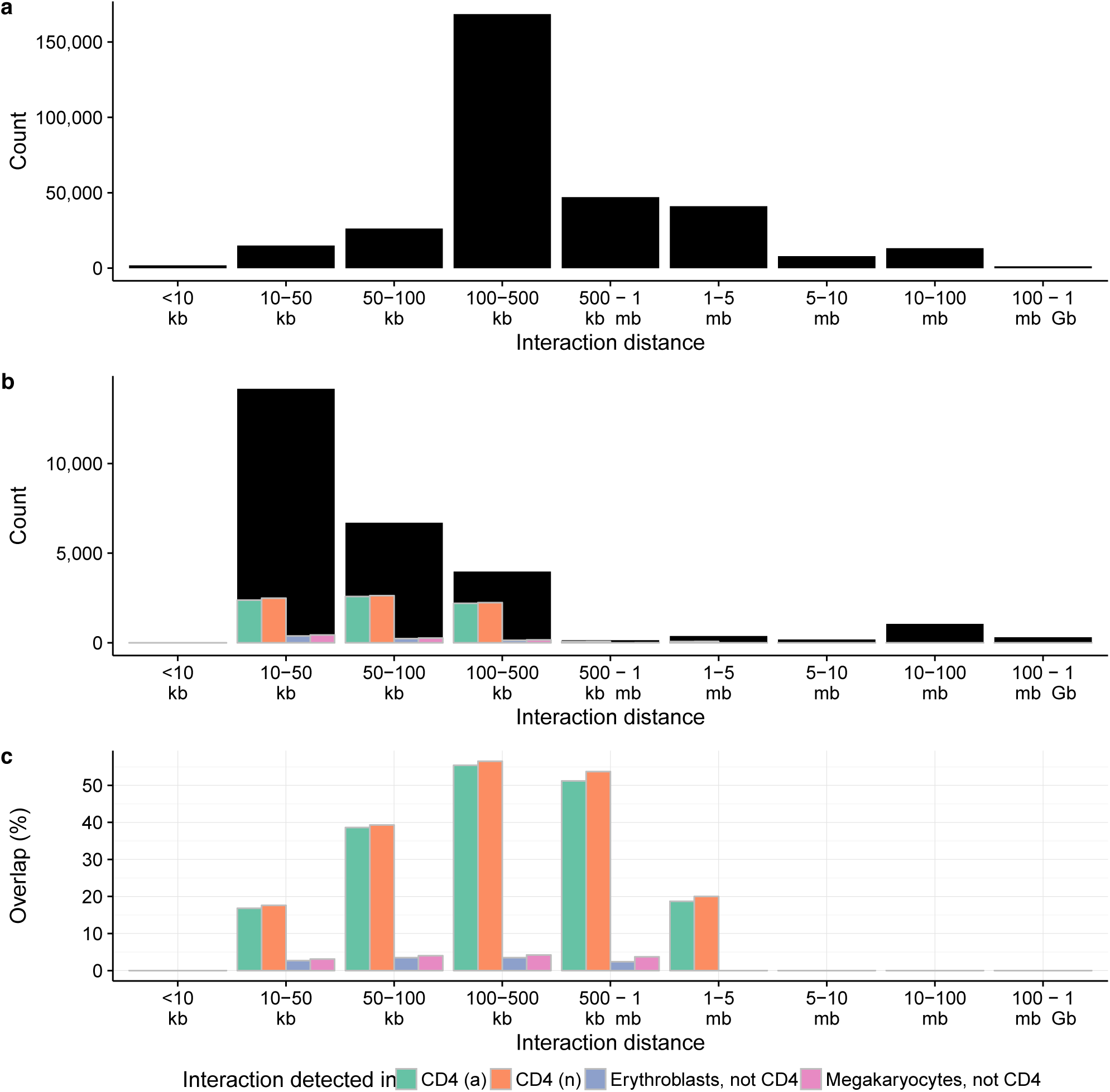
Validation of PCHi-C by ChIA-PET. Distance profiles of PCHi-C and ChIA-PET derived promoter-enhancer interactions in **a** PCHi-C, non-activated CD4^+^ T cells and **b** ChIA-PET (Chepelev et al), black bars. Coloured bars show the count (**b**) or percentage (**c**) of ChIA-PET interactions recovered in the PCHi-C experiment in non-activated and activated CD4^+^ T cells (CD4 (a) and CD4 (n), respectively) and, for comparison, two non-lymphocyte cells, ethryroblasts and megakaryocytes processed in parallel after exclusion of interactions found in either CD4^+^ T cell. Calling interactions requires correction for the expected higher density of random collisions at shorter distances^57^ which are explicitly modelled by CHICAGO^9^ used in this study but not in the ChIA-PET study^12^. As a result, we expected a higher false positive rate from the ChIA-PET data at shorter distances. Indeed, while we replicated only 17% of interactions in the 10-50kb range, we replicated over 50% of the longer range interactions (*>*100 kb).

**Fig. S4:**
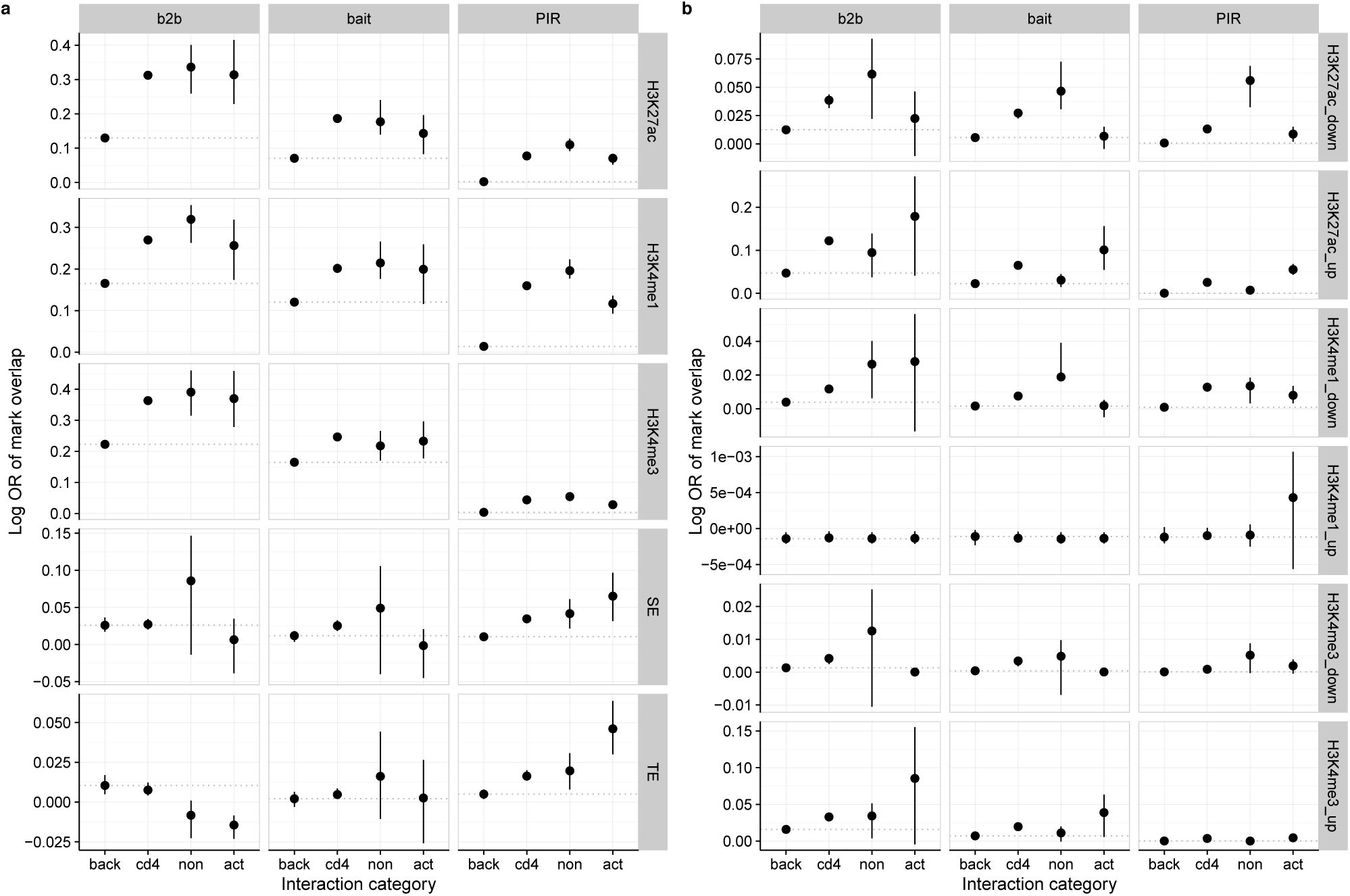
Chromatin state profiles of interacting fragments. Log odds ratio that bait, bait-to-bait (b2b) and PIR regions detected in background cells (back; megakayrocytes and erythroblasts), activated and non-activated CD4^+^ T cells (cd4), specifically non-activated or activated CD4^+^ T cells (non or act, respectively) overlap (**a**) given ChIP-seq peaks or typical (TE) or super (SE) enhancers in resting T cells as previously defined^12^ and (**b**) differential (FDR*<*0.1) ChIP-seq peaks compared to non-interacting regions. Regions considered specific to activated or non-activated cells had a CHICAGO score *>* 5 only in that cell type and were considered differential interactions in a comparative analysis of mapped sequence counts at FDR*<*0.1.

**Fig. S5:**
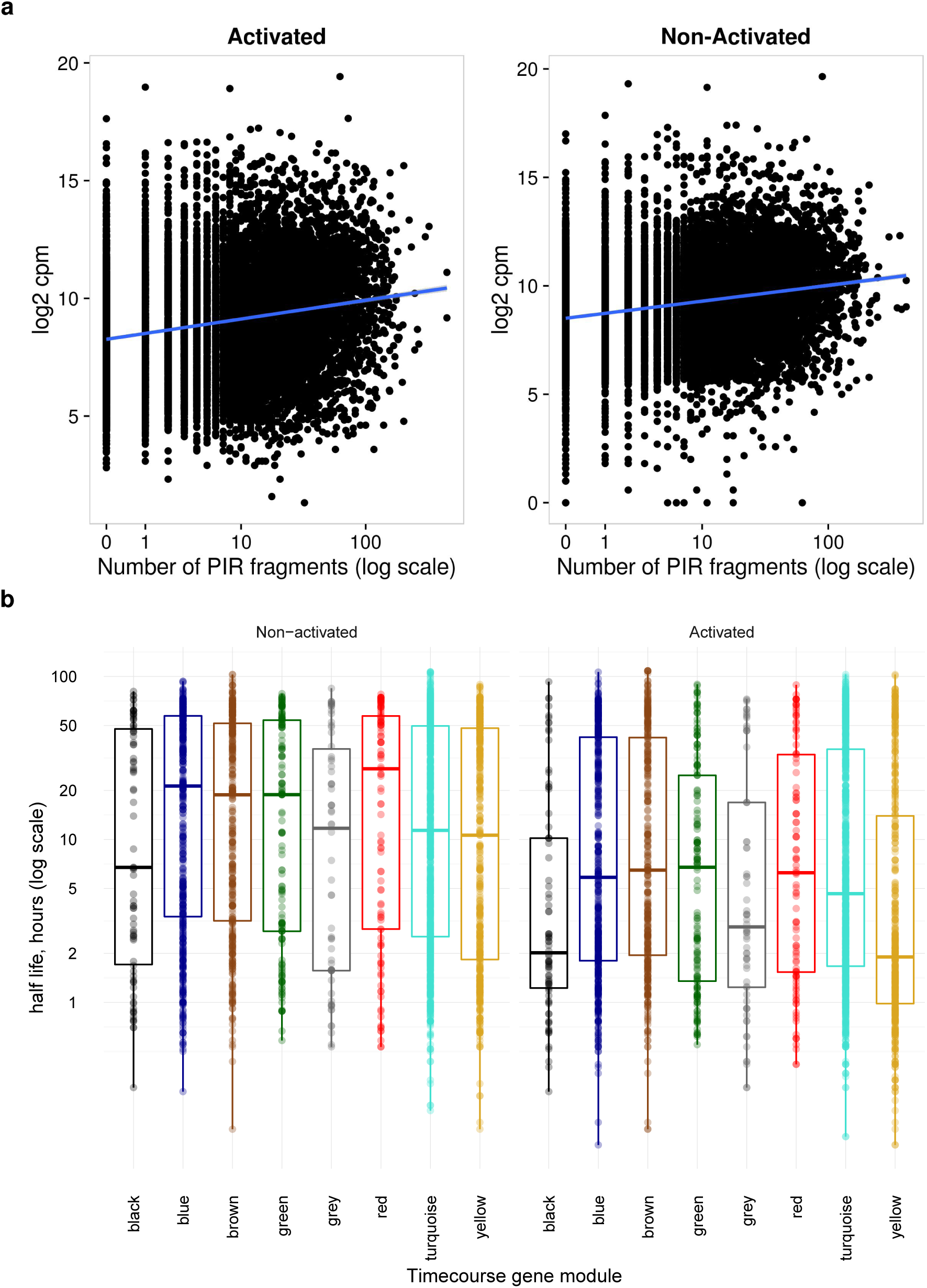
Relationship of gene expression to PIR number and mRNA half-life. **a** RNA-seq expression (counts per million mapped reads, log_2_ scale) shows a positive correlation with the number of PIRs indentified through PCHi-C. **b** half-life of mRNA (Raghavan et al. 2002) by gene module in non-activated and activated cells. The most dynamically regulated genes in our time-course, those in the black module, had the shortest half-life (*p* = 3 *×* 10^*-*8^).

**Fig. S6:**
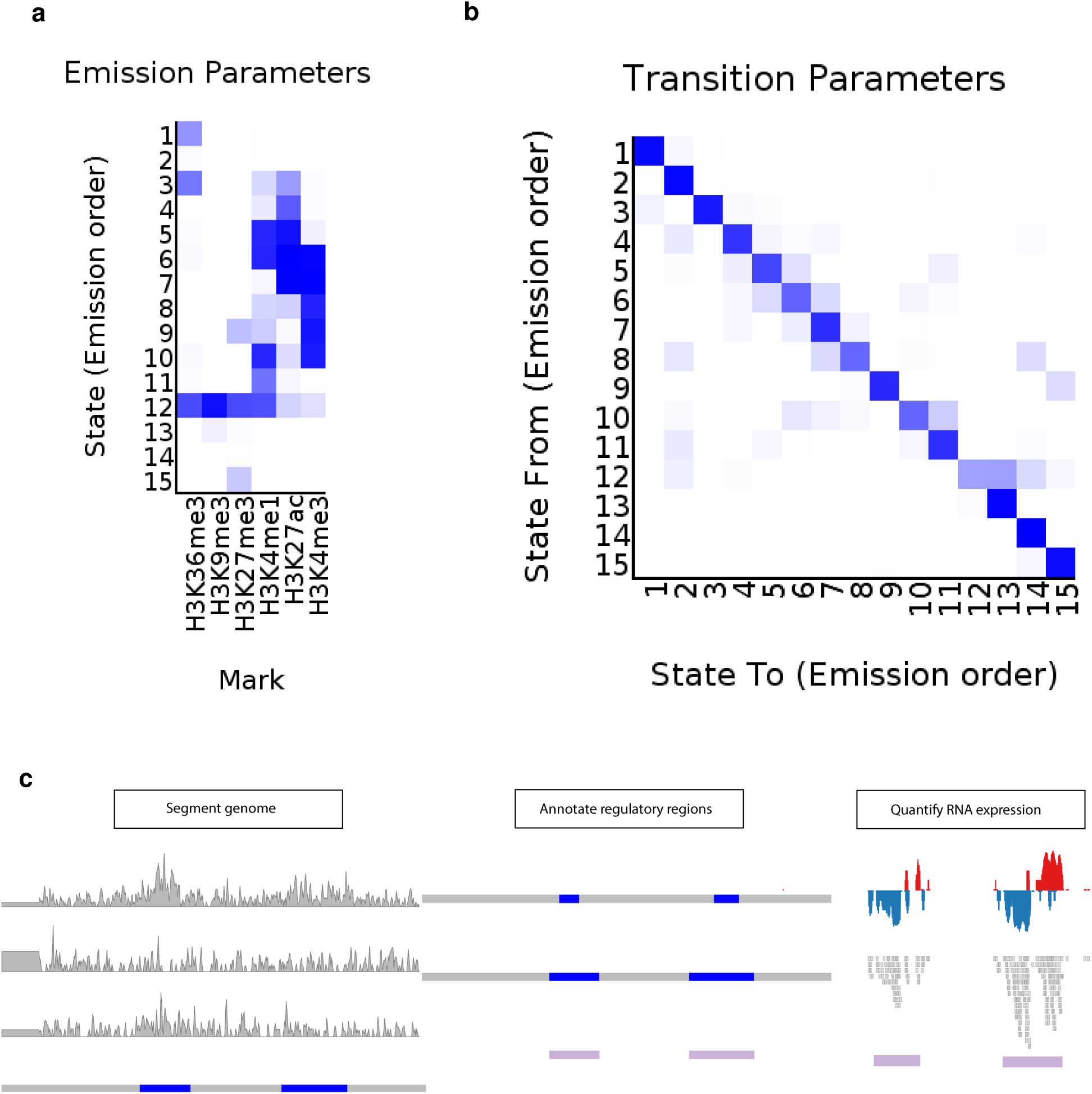
Definition and quantification of regulatory RNAs. CHROMHMM analysis of ChIP-seq marks was used to produce a whole genome segmentation into 15 states. Resulting emission (**a**) and transmission (**b**) matrices are shown. States E4-E11 were defined as regulatory. **c** Neighbouring regions containing promoter or enhancer states (E4-E11) were merged together into regulatory annotations. Expression levels of each regulatory area were quantified using RNA-seq in a strand-aware fashion, to avoid the confounding effect of overlapping genomic features.

**Fig. S7:**
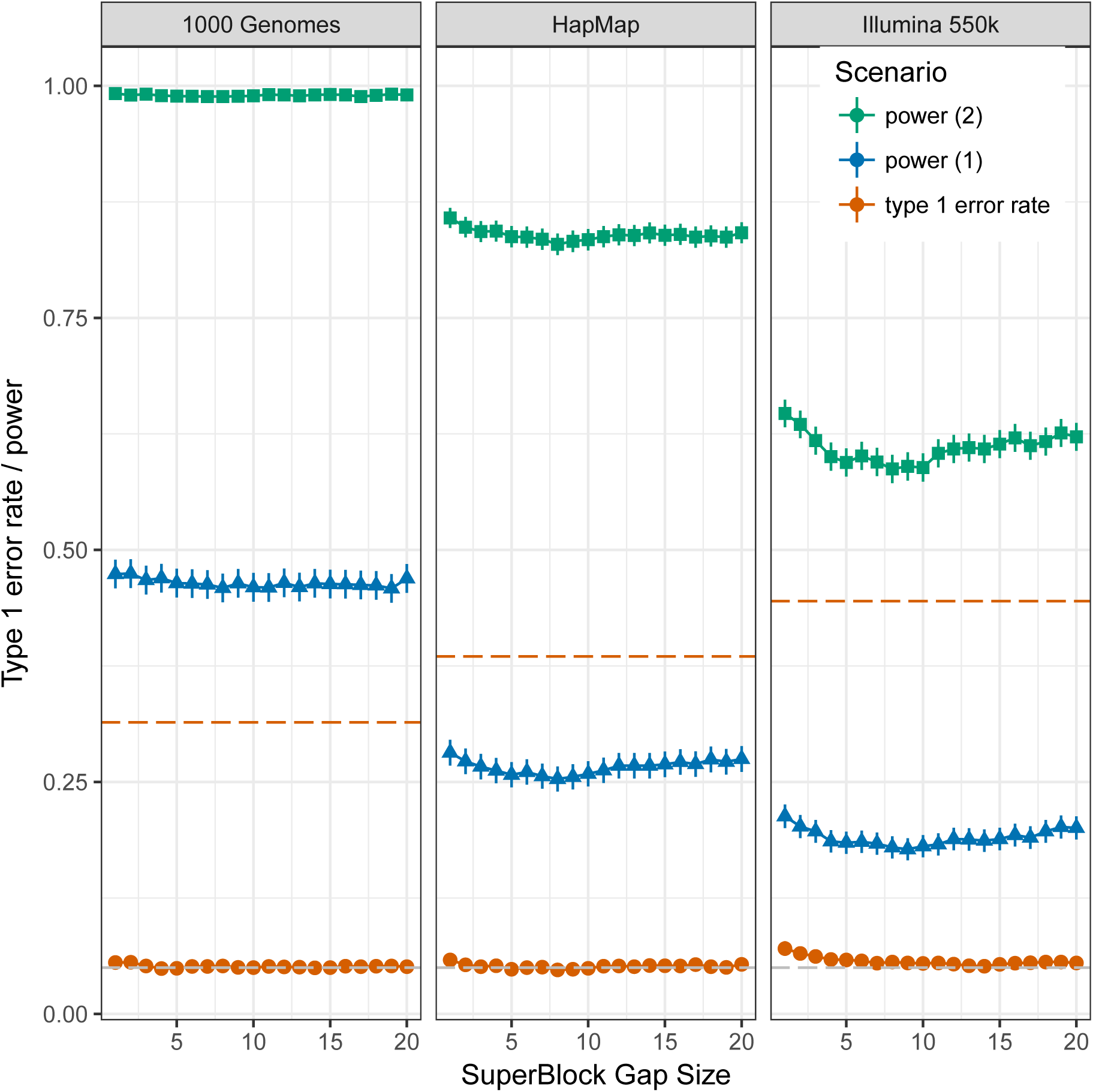
blockshifter calibration. Each panel represents a simulated genotyping density: 1000 genomes (156,082 SNPs); HapMap (44,647 SNPs input,); Ilumina 550k (10,241 SNPs input). Points represent type 1 error rates (alpha=0.05) for the null scenario (no enrichment of GWAS variants in test specific PIRs) and moderate (power 1) and strong (power 2) enrichment scenarios across 4000 simulated GWAS, with differing blockshifter ‘SuperBlock’ gap size parameter. Error bars represent 95% confidence intervals. Dashed red lines represent the type 1 error rate for Fisher’s test of enrichment of variants in test and control PIRs. The naive application of Fisher’s test leads to substantial inflation of type 1 error rate, more so in lower-density genotyping scenarios. Blockshifter maintains type 1 error rate control, although a gap size of 5 or more is required to deal with the extended correlation induced by PMI in lower density genotyping scenarios, while Blockshifter power is impacted, as expected, by genotyping density.

**Fig. S8:**
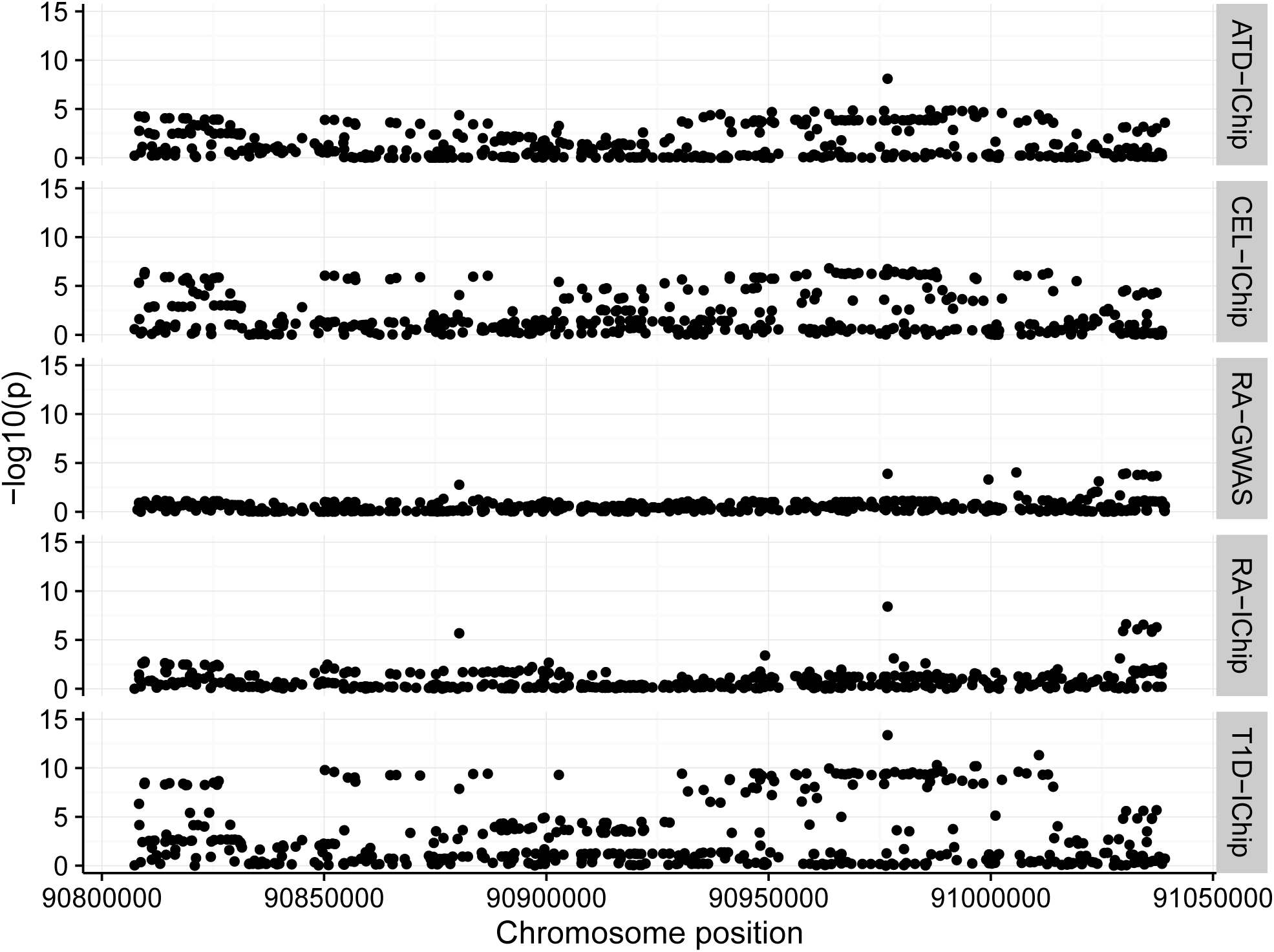
*MDN1* is prioritised for RA through ImmunoChip but not GWAS data. Similar signals are found for ATD and T1D, which also link to *MDN1*, supporting the RA-ImmunoChip result. The lack of prioritisation in the RA-GWAS dataset relates to the weaker evidence for association in this region.

**Fig. S9:**
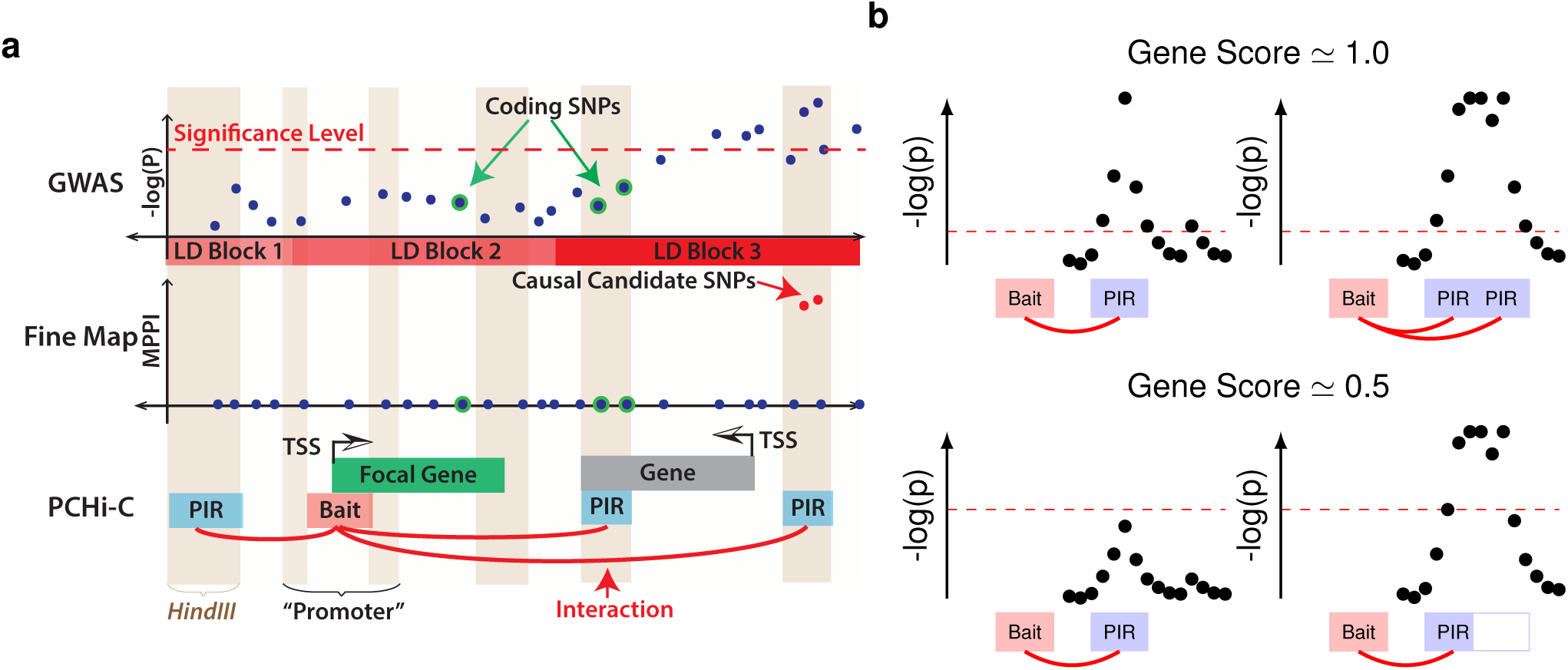
Gene prioritisation using COGS. We prioritised disease candidate causal genes by integrating GWAS data with PCHi-C interactions using the COGS algorithm. **a** The algorithm uses a Bayesian method to define the marginal posterior probability of inclusion (MPPI, middle panel) for each variant from GWAS data (top panel). We can also calculate the MPPI marginalising across PIRs (light blue, bottom panel), coding variants and promoter regions for each focal gene. *Hind*III fragments are indicated by dark/light vertical shading. **b** Note that the gene score is therefore a function of the strength of GWAS signal, how peaked/diffuse it is, and the interactions. For example, in the top row there are two strong GWAS signals, one peaked, one diffuse, but the PIRs cover all of the most strongly associated SNPs, and in each case the gene score is expected to be close to 1. In the lower left plot, the GWAS signal is less strong, not even genomewide significant, but all the most associated SNPs lie within the PIR. The score will fall, perhaps to around 0.5, reflecting the weaker evidence for disease association. In contrast, the bottom left plot shows a diffuse signal, only part of which lies within a PIR. Although we can be confident the disease is genuinely associated, only about half the fine mapped candidate causal SNPs will lie within a PIR, and the gene score will again fall, to about 0.5. The situations in the lower row are quite different, but will generate similar scores.

**Fig. S10:**
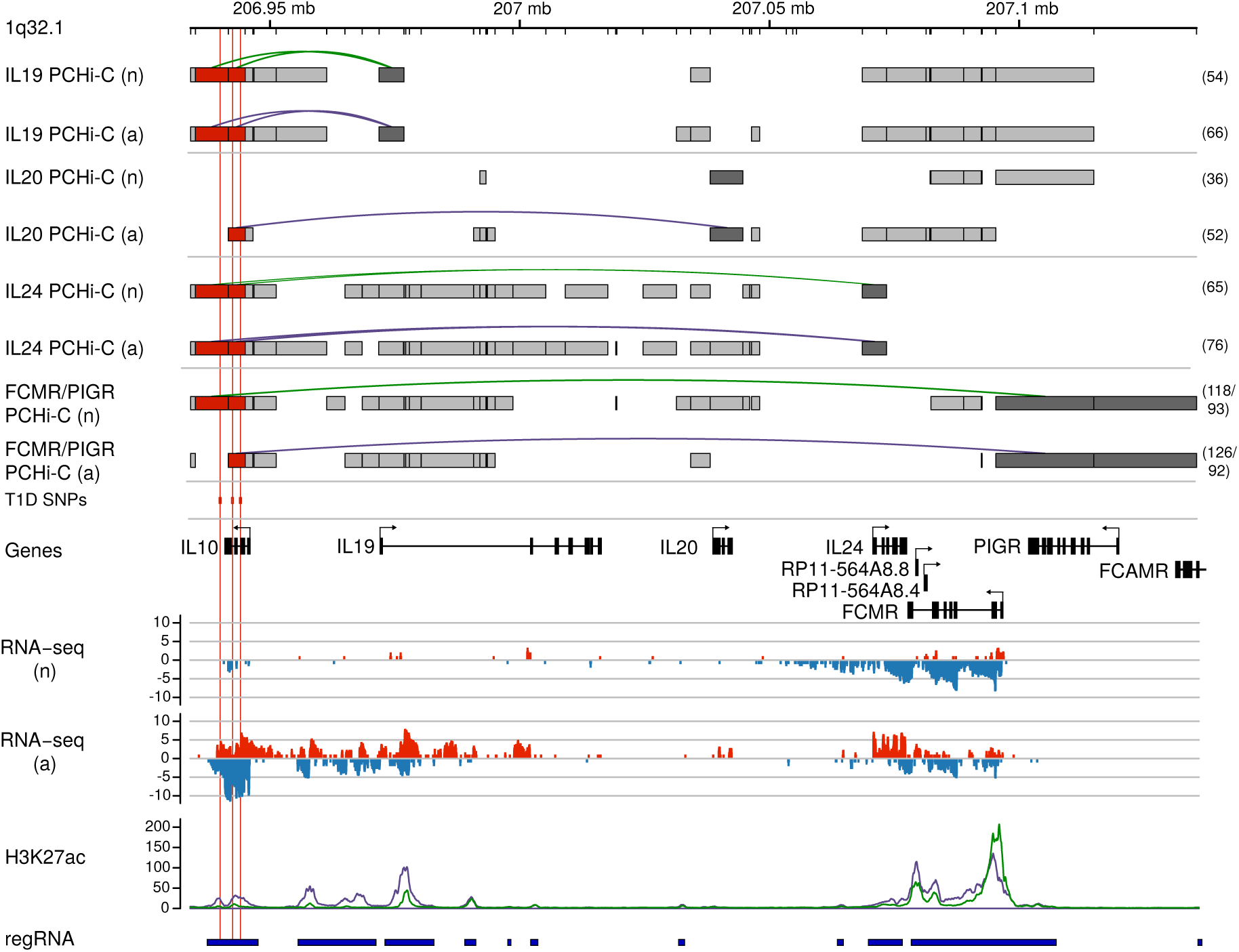
Multiple genes on chromosome 1q32.1 (*IL10*, *IL19*, *IL20*, *IL24*, *FCAMR*/*PIGR*) are prioritised for T1D, CRO and UC. For full legend see **Fig. 5.**

**Fig. S11:**
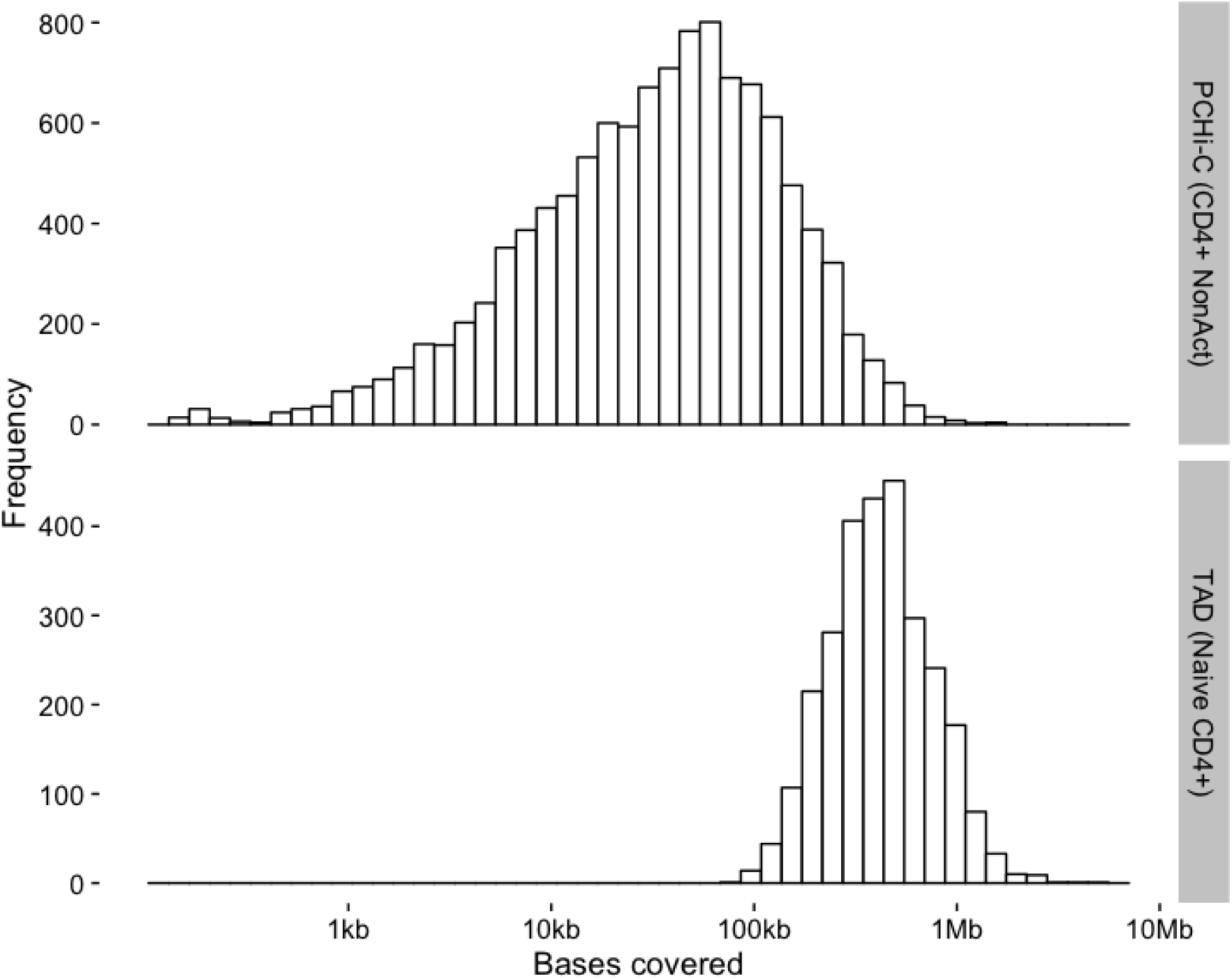
Histograms show the distribution of summed PIR length by gene in non-activated CD4^+^ T cells (top panel) and TAD length in naive CD4^+^ T cells. Note the x axis is drawn using a log scale and that for each gene we have included the promoter-baited fragment and its two immediate neighbours to allow that PCHi-C cannot detect very proximal interactions in this range.

**Fig. S12:**
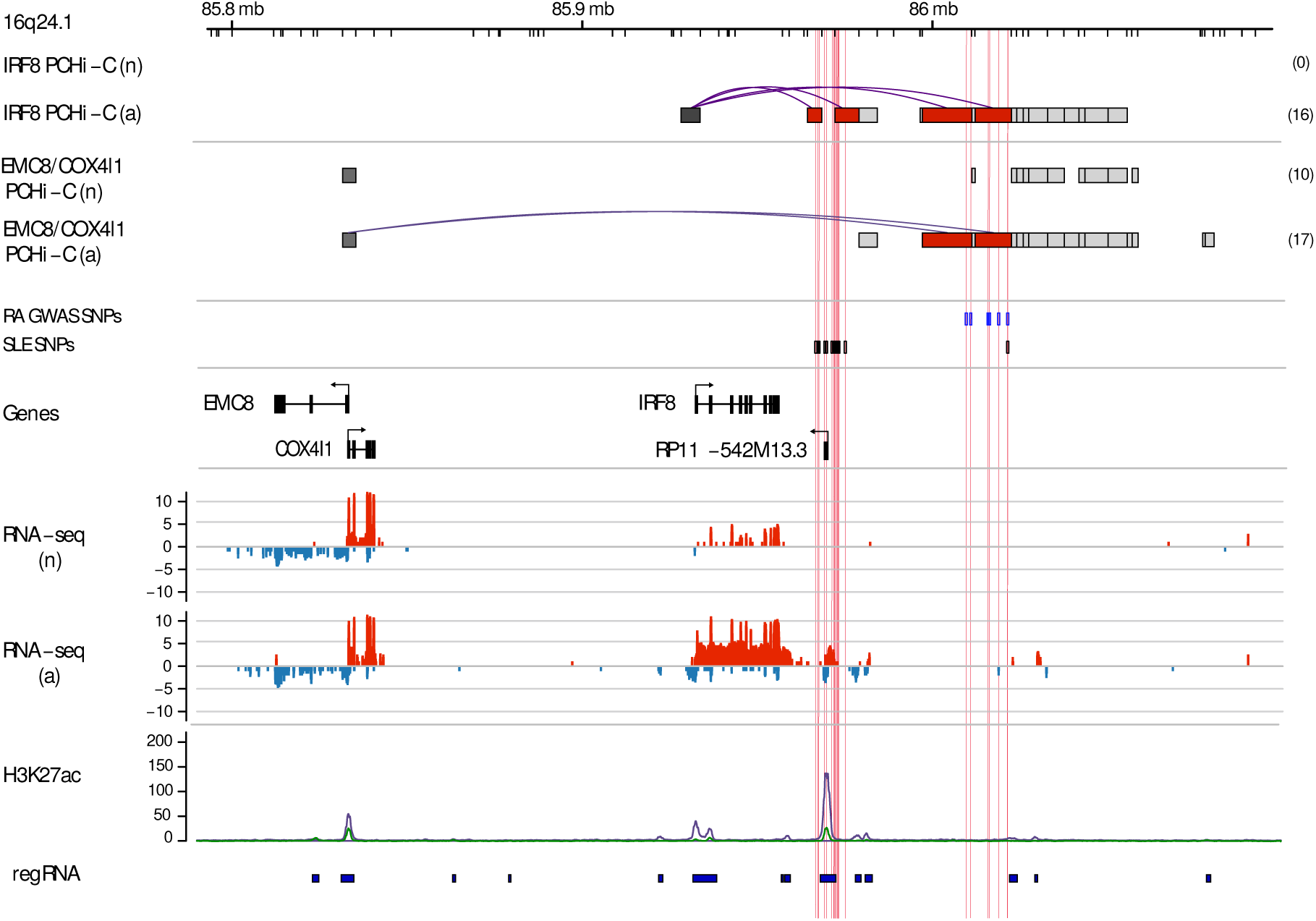
*IRF8* and *EMC8*/*COX4I1* on chromosome 16 are prioritised for RA and SLE. For full legend see **Fig. 5.**

**Fig. S13:**
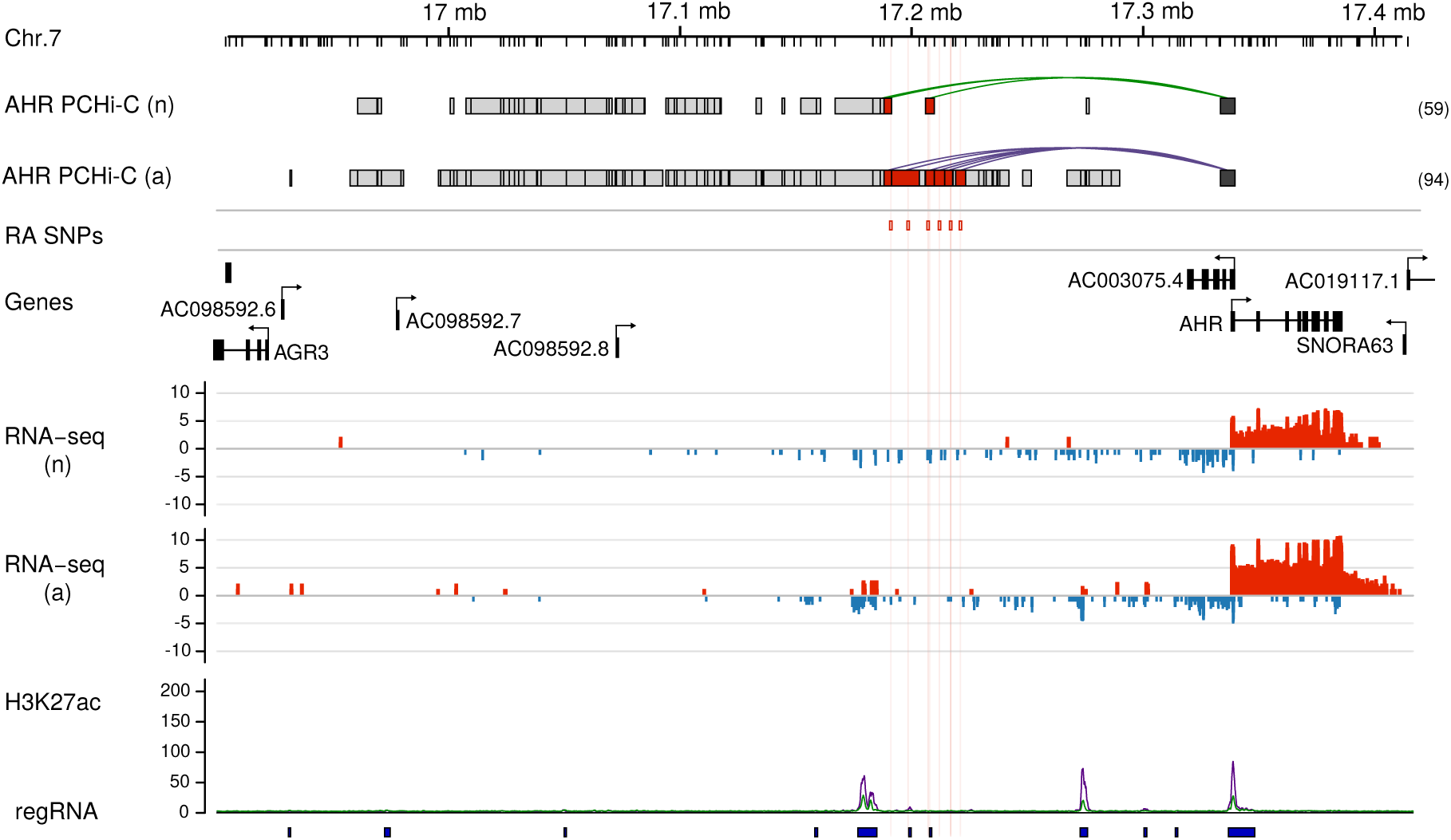
*AHR* on chromosome 7 is prioritised for RA in activated CD4^+^ T cells. For full legend see **Fig. 5.**

**Fig. S14:**
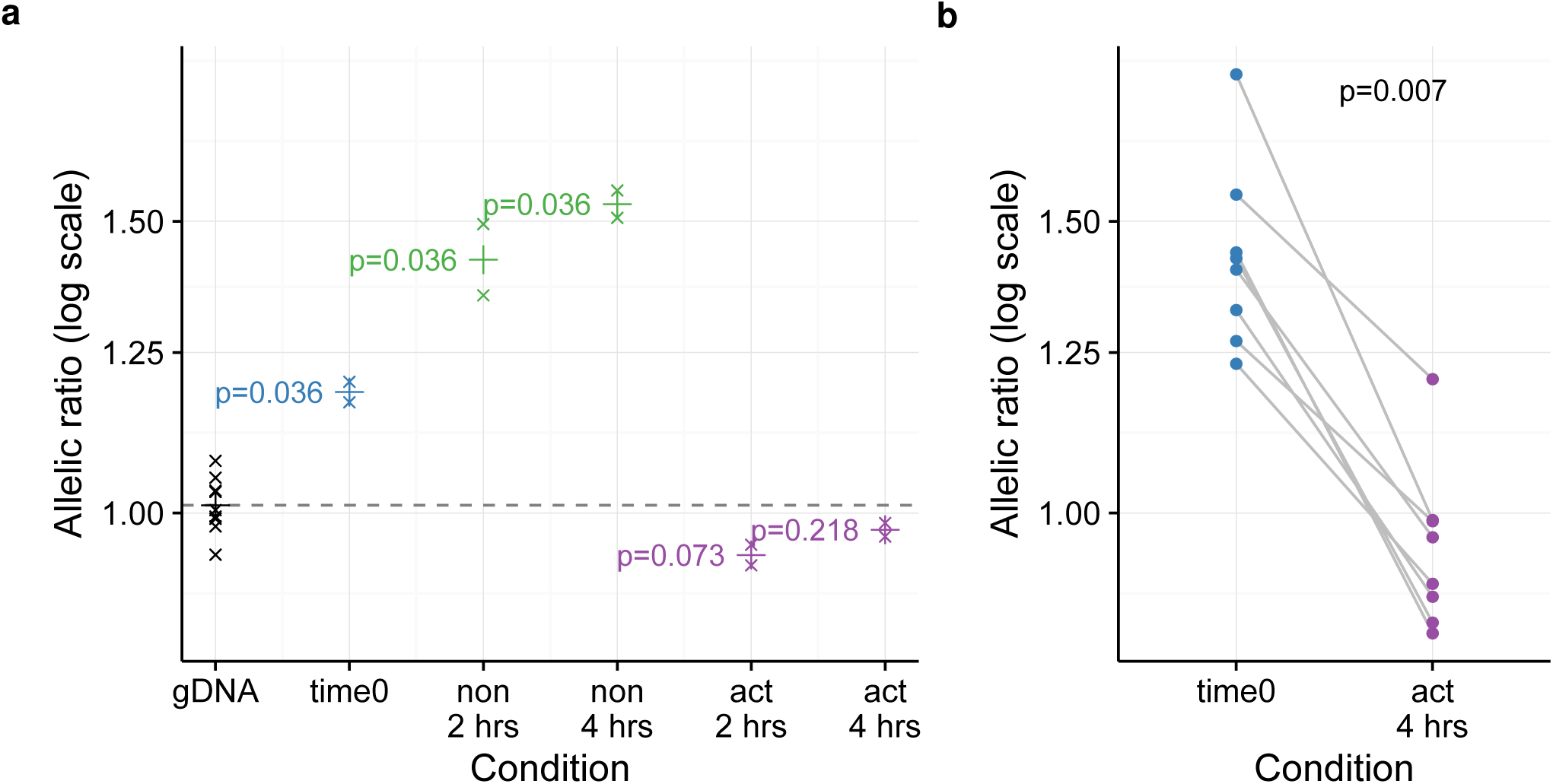
Allelic imbalance in mRNA expression in individuals heterozygous for group A SNPs is confirmed with reporter SNP rs12244380 (*IL2RA* 3’ UTR) **a**: Allelic imbalance in mRNA expression in total CD4^+^ T cells from individuals heterozygous for group A SNPs using rs12244380 as a reporter SNP in non-activated (non) and activated (act) CD4^+^ T cells compared to genomic DNA (gDNA, expected ratio=1). Allelic ratio is defined as the ratio of counts of the allele carried on the chromosome carrying rs12722495:T to that carried on the chromosome carrying rs12722495:C. ‘ *×* ‘=geometric mean of the allelic ratio over 2-3 replicates within each of 4-5 individuals, and p values from a Wilcoxon rank sum test comparing cDNA to gDNA are shown. ‘+’ shows the geometric mean allelic ratio over all individuals. **b**: Allelic imbalance in mRNA expression in memory CD4^+^ T cells differs between *ex vivo* (time 0) and four hour activated samples from eight individuals heterozygous for group A SNPs using rs12244380 as a reporter SNP. p value from a paired Wilcoxon signed rank test is shown.

